# Nanobodies as novel tools to monitor the mitochondrial fission factor Drp1

**DOI:** 10.1101/2023.12.18.572153

**Authors:** Theresa Froehlich, Andreas Jenner, Claudia Cavarischia-Rega, Funmilayo O. Fagbadebo, Yannic Lurz, Desiree I. Frecot, Philipp D. Kaiser, Stefan Nueske, Armin Scholz, Erik Schäffer, Ana J. Garcia-Saez, Boris Macek, Ulrich Rothbauer

## Abstract

In cells, mitochondria undergo constant fusion and fission. An essential factor for fission is the mammalian dynamin-related protein 1 (Drp1). Dysregulation of Drp1 has been linked to neurodegenerative diseases including Parkinson’s as well as cardiovascular diseases and cancer. Here, we developed nanobodies (Nbs) for proteomics, advanced microscopy and live cell imaging of Drp1. To specifically enrich endogenous Drp1 with interacting proteins for proteomics, we functionalized high-affinity Nbs as capture matrices. Furthermore, we detected Drp1 by bivalent Nbs combined with site-directed fluorophore labelling in super-resolution STORM microscopy. For real-time imaging of Drp1, we intracellularly expressed fluorescently labelled Nbs, so-called chromobodies (Cbs). To improve the signal-to-noise ratio, we further converted Cbs into a “turnover-accelerated” format. With these imaging probes, we visualized the dynamics of endogenous Drp1 upon compound-induced mitochondrial fission in living cells. Considering the wide range of research applications, the presented Nb toolset will open up new possibilities for advanced functional studies of Drp1 in disease-relevant models.

## Introduction

Mitochondrial morphology is controlled by the balance between two opposing processes, fusion and fission. These highly regulated events adapt the cellular mitochondrial network to the bioenergetic requirements of the cell and maintain the functional integrity of the mitochondria in important cellular processes such as iron metabolism, lipid biosynthesis, calcium homeostasis and cell death (Iwata et al., 2020; Rasmussen et al., 2020; Vantaggiato et al., 2019). Mitochondrial fission in higher vertebrates is primarily mediated by the large GTPase dynamin-related protein 1 (*DNM1L*; Drp1 also known as DLP1). The fission process follows a coordinated sequence of events in which cytosolic Drp1 is initially recruited to mitochondrial contact sites with the endoplasmic reticulum (ER), facilitated by ER membrane-associated actin filaments (Chakrabarti et al., 2018; Ji et al., 2017; Li et al., 2015). On the mitochondrial outer membrane (MOM), Drp1 associates with adaptor proteins including the mitochondrial dynamics protein of 49 kDa (MiD49, also known as MIEF2), mitochondrial dynamics protein of 51 kDa (MiD51; also known as MIEF1), and presumably the mitochondrial fission factor (MFF) and fission protein 1 (FIS1) (Loson et al., 2013; Osellame et al., 2016; Palmer et al., 2013). Subsequently, Drp1 oligomerizes into ring-like or helical structures surrounding mitochondria and GTP hydrolysis then triggers a conformational change in Drp1 causing the contractive division of the mitochondrial network (Bai et al., 2015; De Vos et al., 2005; Friedman et al., 2011; Frohlich et al., 2013; Korobova et al., 2013). In addition to mitochondria, Drp1 also mediates division of peroxisomes in combination with the adaptor proteins MFF and FIS1(Koch et al., 2003; Koch and Brocard, 2012; Li and Gould, 2002). Dysregulation of Drp1 is associated with neurodegenerative diseases such as Alzheimer’s (Cho et al., 2009; Wang et al., 2009), Huntington’s (Shirendeb et al., 2012; Song et al., 2011) or Parkinson’s (Han et al., 2020) as well as cardiovascular diseases (Jin et al., 2021; Nan et al., 2017). In addition, it has been linked to cancer, including melanoma, glioblastoma, lung, breast, thyroid, pancreatic and head and neck cancer (Ferreira-da-Silva et al., 2015; Huang et al., 2022; Kashatus et al., 2015; Liang et al., 2020; Rehman et al., 2012; Serasinghe et al., 2015; Zhao et al., 2013). These findings highlight the growing importance of Drp1 as a biomarker or potential target for therapeutic interventions. However, despite several *in vitro* and *in vivo* studies, detailed information on the molecular mechanism and structural changes of Drp1 during mitochondrial fission, as well as its cellular dynamics and interacting components, is still lacking (Giacomello et al., 2020; Tong et al., 2020; Zerihun et al., 2023). This gap in knowledge is partly due to the limited availability of research tools to study Drp1. Notably, most live cell analyses rely on ectopic expression of fluorescent fusion constructs or epitope-tagged Drp1 (Ji et al., 2017; Labrousse et al., 1999; Michalska et al., 2018; Solesio et al., 2013; Xiong et al., 2022). However, recent studies revealed that N- or C-terminal labelling of Drp1 with fluorescent proteins or epitope tags leads to altered oligomerization dynamics and impairs its GTPase activity (Montecinos-Franjola et al., 2020).

Single-domain antibody fragments derived from heavy chain-only antibodies of camelids (Hamers-Casterman et al., 1993), also known as nanobodies (Nbs), have emerged as versatile research tools in biomedical research. Due to their specific binding properties, small size, high stability and good solubility, these binders became an attractive alternative to conventional antibodies in many biochemical and cell biological research applications (Frecot et al., 2023). In addition, intracellular functional Nbs genetically fused to fluorescent proteins, so-called chromobodies (Cbs), serve as imaging probes to visualize dynamic changes of target antigens without genetic or covalent modifications and can be applied in different cellular compartments as well as in whole organisms (Wagner and Rothbauer, 2020). Here, we have identified a set of novel Nbs specifically recognizing human Drp1. Following their detailed biochemical, biophysical and functional characterization, we have developed them as broadly applicable research tools. Thus, we converted them into affinity matrices for proteomic analysis of Drp1 and developed bivalent Nbs as labelling probes to detect endogenous Drp1 in confocal and super-resolution microscopy (SRM). By engineering turnover-accelerated Drp1 Cbs, which have an improved signal-to-noise ratio, we were able to visualize the drug-induced recruitment of Drp1 to mitochondria in living cells. Considering the broad spectrum of applications, we propose that the Drp1-specific Nbs/Cbs described herein open up new possibilities for future advanced functional studies of Drp1 in disease-relevant models and represent a promising alternative to currently available approaches.

## Results

### Identification of Drp1-specific Nbs

To generate Nbs against human Drp1 (Drp1), we immunized an alpaca (*Vicugna pacos*) with recombinant Drp1 using a 91-day immunization protocol and detected a specific immune response in a serum ELISA 63 days after the first vaccination (**Fig. S1A**). Subsequently, we established a Nb phagemid library (size ∼ 2 x 10^7^ clones) from the mRNA of the peripheral B lymphocytes, which was subjected to phage display using either passively adsorbed or biotinylated Drp1. After two rounds of biopanning, we analysed 260 individual clones in a solid-phase phage ELISA and identified 27 positive binders (**Fig. S1B**) with eight unique sequences (**Table S1**). Notably, only two Nbs, D7 and D63, displaying highly diverse complementarity determining regions 3 (CDR3) (**Fig. 1A**), showed sufficient binding signals in a protein ELISA using periplasmic extracts derived from Nb-expressing *Escherichia coli* (*E. coli*) (**Fig. S1C**). Therefore, we purified these binders by immobilized metal ion affinity chromatography (IMAC) and subsequent size exclusion chromatography (SEC) (**Fig. 1B**). For determining their binding affinities, both purified Nbs were subjected to biolayer interferometry (BLI). Biotinylated Nbs were immobilized on streptavidin (SA) biosensors and binding kinetics were measured by titrating varying concentrations of Drp1. Both Nbs, D7 and D63, showed affinities in the low nanomolar range with K_D_ values of ∼3.5 nM and ∼1.8 nM, respectively (**Fig. 1C**, **Fig. S2**). In addition, we analysed their folding stability using nano-differential scanning fluorimetry (nanoDSF). This revealed that both Nbs exhibit stable folding and a low tendency to aggregate having melting temperatures of ∼68°C and ∼70°C for D7 and D63, respectively (**Fig. 1D**). To further assess whether Nb binding affects the functionality of Drp1, we analysed the GTPase activity of Drp1 in the presence of the respective Nbs *in vitro*. While D7 and the negative control (GFP Nb) had no effect on the enzymatic activity of Drp1, we observed a significant increase in the GTPase rate by a factor of ∼ 1.7 after addition of a 10-fold molar excess of D63 (**Fig. 1E**).

**Figure 1:**
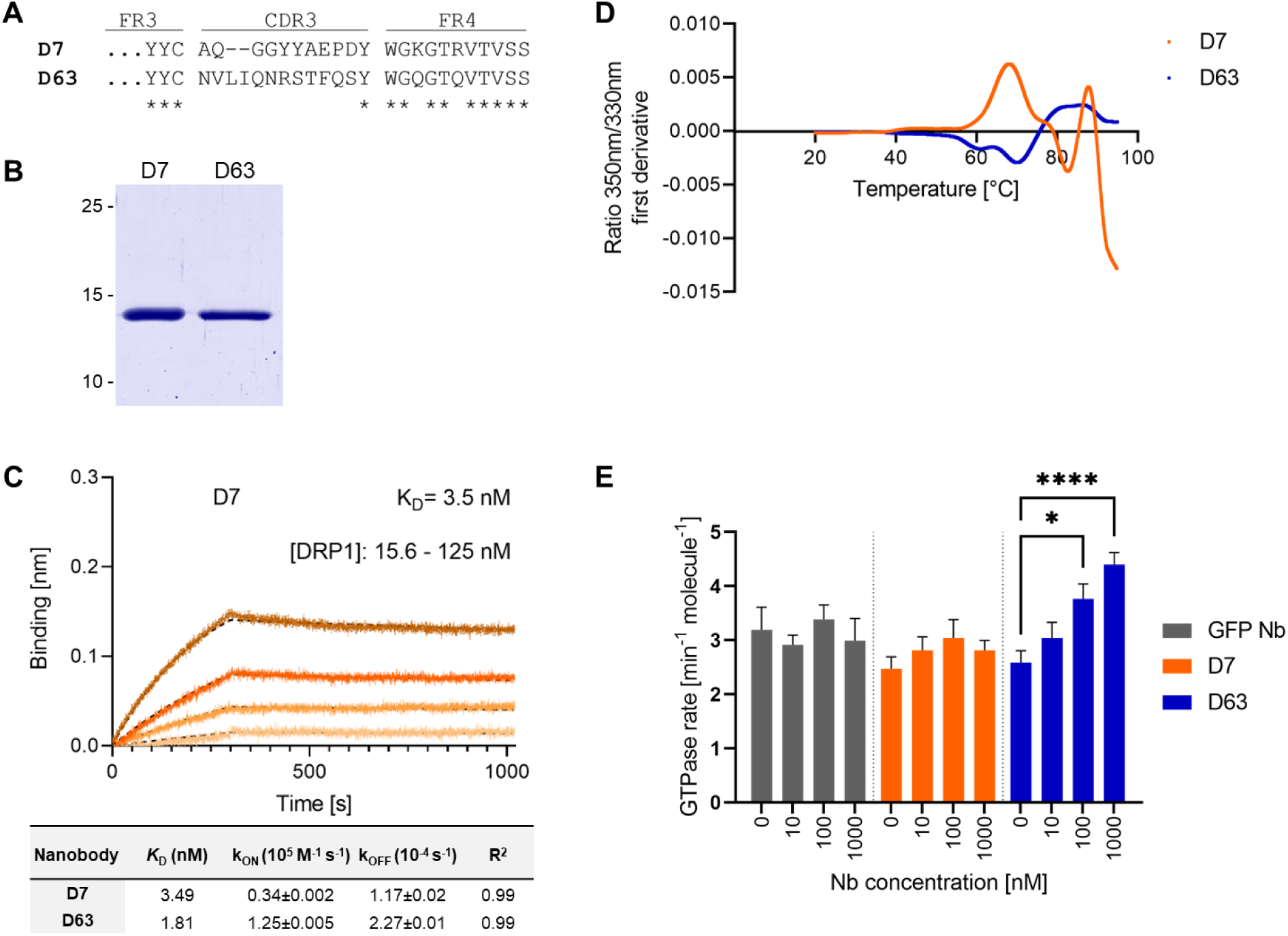
Identification and biochemical characterization of Drp1-specific Nbs. (**A**) Amino acid (aa) sequences of the complementary determining region 3 (CDR3) of the Nbs D7 and D63 identified by protein ELISA. (**B**) Recombinant expression and purification of D7 and D63 by immobilized metal affinity chromatography (IMAC) and size exclusion chromatography (SEC). Coomassie staining of purified Nbs (2 µg) is shown (**C**) Biolayer interferometry (BLI)-based affinity measurements exemplarily shown for D7. Biotinylated Nb D7 was immobilized on streptavidin sensors. Kinetic measurements were performed using four concentrations of purified Drp1 ranging from 15.6 – 125 nM (displayed with gradually darker shades of color). The binding affinity (K_D_) was calculated from global 1:1 fit shown as dashed lines. Affinities (K_D_), association constants (*k*_on_) and dissociation constants (*k*_off_) of D7 and D63 determined by BLI shown as mean ± SD. (**D**) Stability analysis by nano-differential scanning fluorimetry (nanoDSF) displaying the fluorescence ratio (350nm/330nm) first derivative for D7 (orange) and D63 (blue). Data are shown as the mean value of three technical replicates. (**E**) GTP hydrolysis of 100 nM Drp1 in solution in the presence of increasing concentrations of different Nbs at 37°C (mean ± SED, n = 5 experiments on 4 days). Significance was tested using two-way ANOVA and multiple comparison analysis in GraphPad Prism (n.s. p > 0.05, * p < 0.05, **** p < 0.0001).

### Selected Drp1-Nbs recognize different domains of Drp1 and precipitate endogenous Drp1

Considering that Drp1 is composed of multiple domains including an N-terminal GTPase (aa 1-337), an unfolded (middle, stalk, aa 338-502), a variable (B insert, aa 503-635), and a C-terminal GTPase effector domain (GED) region (aa 636-736) (Otera et al., 2013), we constructed a series of GFP-labelled domain deletions of Drp1 (**Fig. 2A**) and tested whether the selected Nbs recognize specific domains within Drp1 by pull-down assays. To this end, we generated so-called Drp1 nanotraps for which we covalently immobilized D7 and D63 on N-hydroxysuccinimide (NHS)-activated Sepharose beads via primary amino groups of accessible lysine residues (Fagbadebo et al., 2022; Traenkle et al., 2020). For binding analysis, the Drp1 nanotraps were incubated with soluble protein fractions of HEK293 cells transiently expressing the Drp1 deletion constructs. Here we used the GFP-Trap as a positive control (Rothbauer et al., 2008). Immunoblot analysis revealed that both Nbs precipitated full-length Drp1 as expected (**Fig. 2 B**). However, while D7 exclusively recognizes the N-terminal GTPase domain, D63 only binds in the presence of the GED, whereas none of the Nbs binds non-specifically to GFP (**Fig. 2B**). Next, we tested the capacity of the Drp1 nanotraps to capture endogenous Drp1. Therefore, we generated soluble protein extracts from three different human cell lines (HEK293, HeLa, and U2OS) and incubated them with the Drp1 nanotraps. For comparison, we used a monoclonal anti-Drp1 antibody immobilized on proteinA/G-Sepharose as positive control (PC) and the GFP-Trap as negative control (NC). Immunoblot analysis of all cell lines tested showed that both Drp1 nanotraps precipitated higher levels of endogenous Drp1 compared to the conventional antibody (**Fig. 3A** and **Fig. S3**).

**Figure 2:**
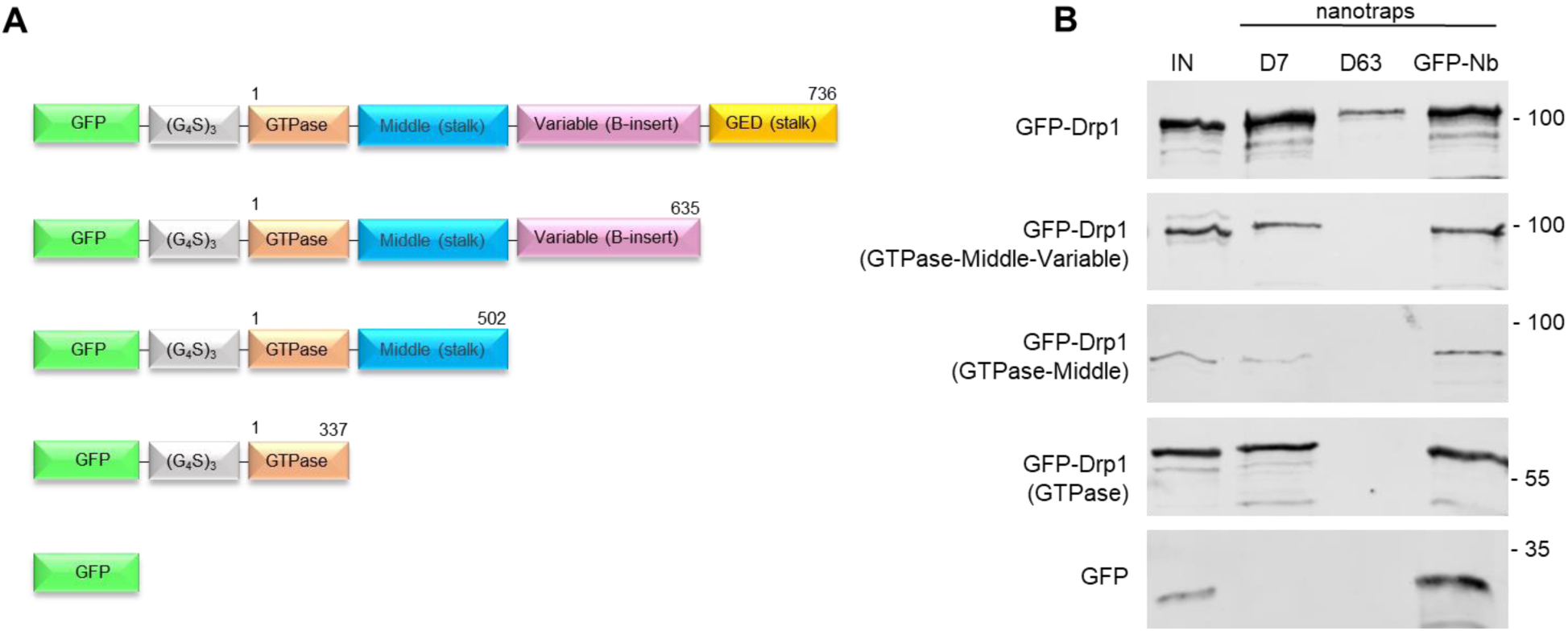
Domain mapping of Drp1-Nbs. (**A**) Schematic illustration of the GFP-labelled Drp1 deletion constructs used for domain binding studies. Numbers indicate amino acid positions of the Drp1 coding sequence (**B**) Results of the pulldown analysis of the individual Drp1 deletions as depicted in (A) using the Drp1-specific nanotraps (D7, D63) or the GFP-Nb as positive control. Input (IN, 1 % of total) and bound (20 % of total) fractions were subjected to SDS-PAGE followed by immunoblot analysis using an anti-GFP antibody. Molecular weights in kDa are indicated on the right.

**Figure 3:**
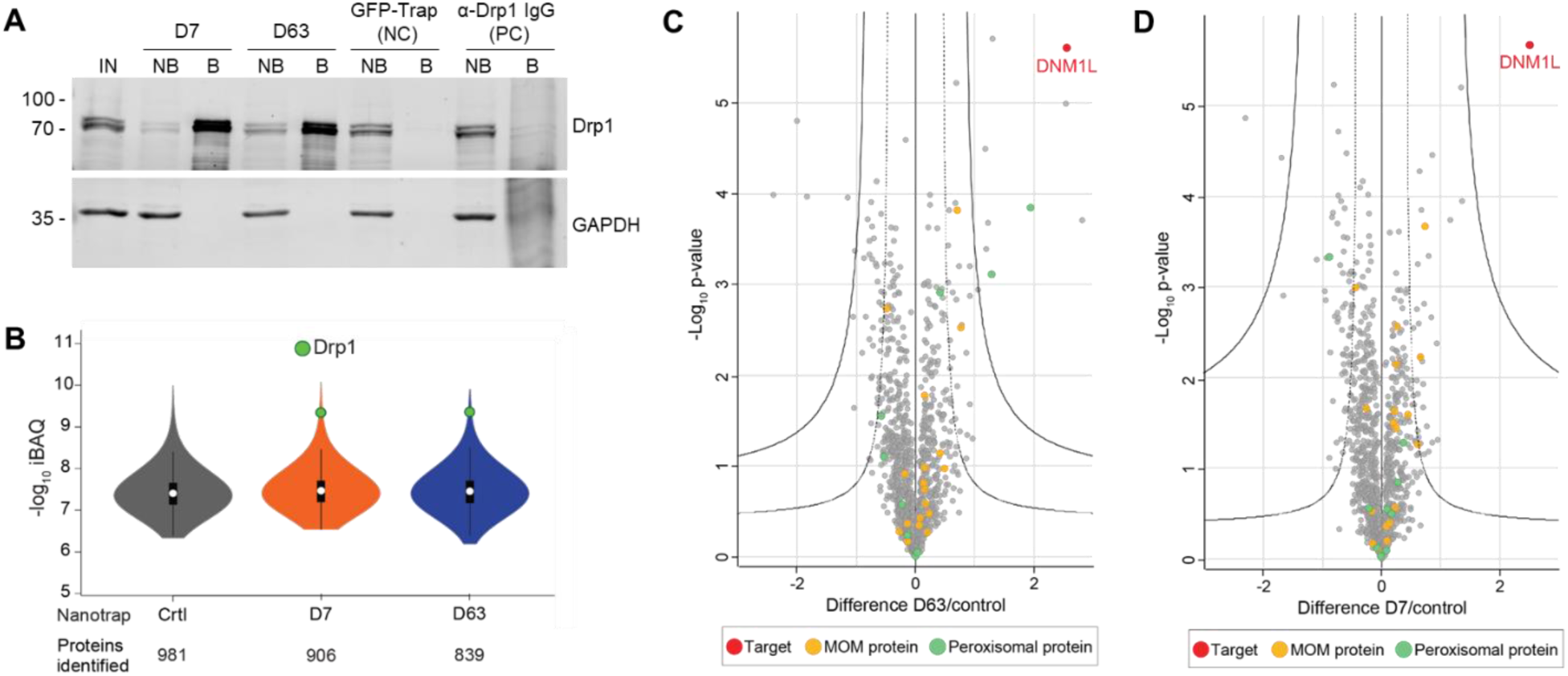
Immunoprecipitation of endogenous Drp1 with mass spectrometry analysis of the interactome of the Drp1-nanotraps. (**A**) Representative result of an immunoprecipitation of endogenous Drp1 with Drp1-specific nanotraps (D7 and D63) using HeLa cells. As negative control (NC), the GFP-Trap as non-specific nanotrap was used and bead-coupled anti-Drp1 IgG served as a positive control (PC). Input (IN, 1 % of total), non-bound (NB, 1 % of total) and bound (B, 33 % of total) fractions were subjected to SDS-PAGE followed by immunoblot analysis using antibodies specific for Drp1 (upper panel) and GAPDH (lower panel). Molecular weights in kDa are indicated on the left. (**B**) Violin plot of the averaged iBAQ intensities of the proteins identified from immunoprecipitation using HEK cells. White circles show the median; box limits indicate the 25^th^ and 75^th^ percentiles as determined by R software; whiskers extend 1.5 times the interquartile range from the 25^th^ and 75^th^ percentiles; polygons represent density estimates of data and extend to extreme values. Drp1 is marked in green. Multi-volcano analysis (Hawaiian plot) of the pull down of D7 (**C**) and D63 (**D**) against the negative control. Log2 transformed ratios of the difference between the Drp1-nanotrap and the control (x axis) are plotted against the log10 transformed p-value (y axis). Drp1 is marked in red, MOM proteins are marked in yellow and peroxisomal proteins in green. Significant interactors can be class A hits (higher confidence, *s*_0_ = 0.1, FDR = 0.01) or class B hits (lower confidence, *s*_0_ = 0.1, FDR = 0.05), thresholds are displayed as a solid line and a dashed line respectively.

For a more detailed insight, we analysed the proteome of both Drp1 nanotraps by mass spectrometry (MS). Three replicates were performed, each with the same starting number of HEK293 cells as the input material for each nanotrap. Overall, the results were highly reproducible showing a Spearman rank correlation close to one (**Fig. S4A**). As expected, the correlation between the two different Drp1 nanotraps was higher than between each nanotrap and the control (GFP-Trap), as confirmed by principal component analysis (PCA) (**Fig. S4B**). We next evaluated how well each nanotrap captured endogenous Drp1. Both nanotraps enabled the identification of multiple Drp1 peptides comparably, resulting in high coverage of the Drp1 sequence (∼70%) (**Fig. S4C**). Notably, 55 “Razor” peptides matching Drp1 isoform 1, 5 or 7 were found in the precipitates of both nanotraps (**Fig. S4D**). However, based on our dataset, we could not distinguish between these isoforms. Next, we confirmed the efficacy of Drp1 enrichment. Overall, the number of protein groups (PGs) identified was comparable for all samples, with a background of up to 981 proteins. For both nanotraps, Drp1 was one of the most abundant proteins detected, while it was not identified in the negative control (**Fig. 3B**). Furthermore, Drp1 was the most significantly enriched protein with both traps (**Fig. 3C,D**). Finally, we scanned our MS data for potential interaction partners of Drp1. Interestingly, both Drp1 nanotraps facilitated the enrichment of cytoplasmic or mitochondrial interactors compared to the control, while only the D63 nanotraps allowed for enrichment of class A interactors of peroxisomal proteins such as HSDL2 (hydroxysteroid dehydrogenase-like protein 2) or HSD17B4 (peroxisomal multifunctional enzyme type 2) (**Fig. 3C,D; Table S2 and Table S3**). In addition, with both Drp1 nanotraps, we found a significant enrichment of CDGSH iron-sulfur domain-containing protein 1 (CISD1) and Histone deacetylase 6 (HDAC6), two factors previously reported to interact with Drp1 (English and Barton, 2021; Hua et al., 2021) (**Table S2 and Table S3**). In summary, these results showed that the selected Nbs recognize different domains of Drp1 and can be readily converted into functional capture reagents to precipitate endogenous Drp1 together with its interaction partners.

### bivD7 enables immunofluorescence detection of endogenous Drp1

To further examine the potential of the Drp1 Nbs as research tools, we investigated their performance in immunofluorescence (IF) microscopy. First, we applied D7 and D63 as primary binding molecules in combination with a fluorescently labelled anti-VHH antibody in fixed and permeabilized U2OS cells transiently expressing GFP-Drp1. Fluorescence images showed cytosolic colocalization with GFP-Drp1 signal only for D7, whereas D63 showed no staining (**Fig. S5**). However, when we tested staining of endogenous Drp1, we did not observe any specific staining or colocalization with signals from a conventional Drp1 antibody (**Fig. 4A**). Considering that a bivalent format of Nbs can be superior to monovalent versions for imaging purposes (Fagbadebo et al., 2022; Virant et al., 2018), we genetically fused two D7 or two D63 Nbs head-to-tail connected by a flexible Gly-Ser linker [(G_4_S)_4_] and generated a bivalent format of D7 (bivD7) or of D63 (bivD63), respectively. Additionally, we inserted a sortase-tag for site-directed labelling. Both bivalent Nbs were purified as secreted proteins from ExpiCHO cells (**Fig. S6A**) followed by measuring their binding affinities as described for the monovalent versions. The results showed that bivD7 had an increased affinity in the subnanomolar range due to decelerated dissociation, whereas the affinity of bivD63 did not increase compared to the monovalent version (**Fig. S6B,C**). Initially, we tested both bivalent Nbs for colocalization with GFP-Drp1 in U2OS cells and observed more specific signals compared to the monovalent versions, with bivD7 tending to recognize Drp1 more potently compared to bivD63 (**Fig. S7**). Next, we applied bivD7 and bivD63 for staining of endogenous Drp1. Notably, only the bivD7 showed a clear colocalization with the antibody-labelled Drp1 in U2OS cells (**Fig. 4B**). To further confirm that bivD7 specifically recognizes Drp1, we performed immunofluorescence (IF) imaging of wild type (wt) and Drp1 knock-out (KO) HeLa cells. While a colocalization with the Drp1 antibody signal was detected in wt HeLa cells, no Nb-derived signal was observed in Drp1 KO HeLa cells (**Fig. 4C, Fig. S8**). From these results, we concluded that the bivalent format of D7 is a suitable probe for IF applications to visualize endogenous Drp1.

**Figure 4:**
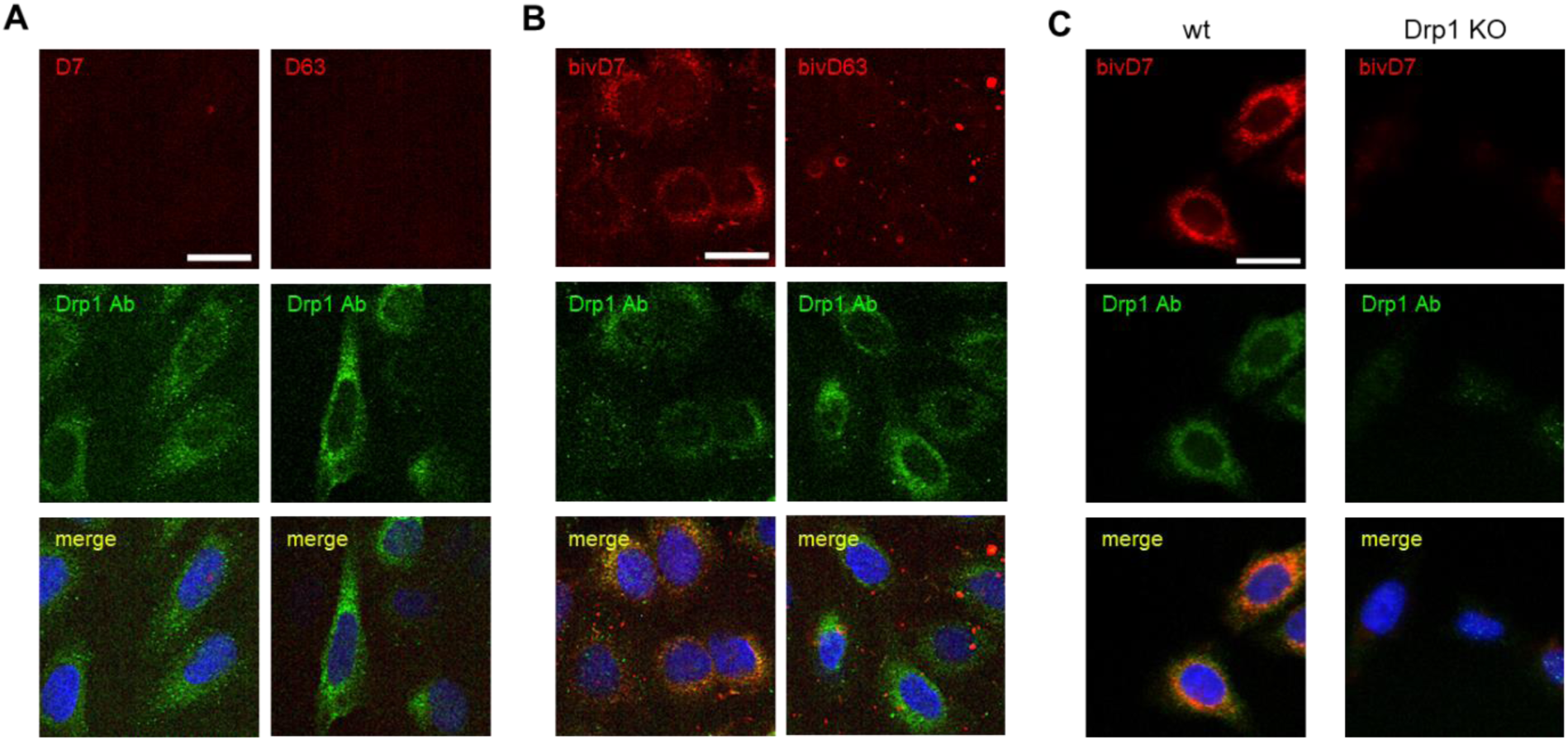
Immunofluorescence staining with Drp1-Nbs. (**A**) Confocal laser scanning microscopy (CLSM) images of U2OS cells stained with D7 and D63 Nbs as primary labelling probes detected with an anti-VHH antibody labelled with Cy5. As positive control, cells were co-stained with anti-Drp1 antibody (Drp1 Ab) followed by detection with an AlexaFluor488-labelled secondary antibody. (**B**) Immunofluorescence (IF) detection of Drp1 in fixed and permeabilized U2OS cells after staining with bivalent versions of D7 and D63 (bivD7, bivD63) as described in (A). (**C**) Immunofluorescence (IF) detection of Drp1 in fixed and permeabilized wt HeLa (left column) and Drp1 KO HeLa cells (right column) stained with bivD7 as described in (B). (**A-C**) Representative images from three independent biological replicates are shown. Nuclei were counterstained with DAPI. Scale bar 25 µm.

### bivD7 is suitable for super-resolution microscopy (SRM)

Due to their small size and their ability to access dense cell compartments and structures, Nbs have been previously described as highly versatile probes for SRM (Cramer et al., 2019; Driouchi et al., 2022; Götzke et al., 2019; Koch and Brocard, 2012; Maidorn et al., 2019; Virant et al., 2018). Hence, we tested the bivD7 in SRM using stochastic optical reconstruction microscopy (STORM). For site-directed labelling, we introduced a peptide comprising an azide group at the C-terminus by chemoenzymatic sortagging (Popp and Ploegh, 2011; Virant et al., 2018), followed by addition of a dibenzocyclooctyne (DBCO) derivative by click chemistry (Fagbadebo et al., 2022) which allowed us to flexibly and specifically conjugate STORM-compatible dyes such as AlexaFluor 647 (AF647). For initial validation, we stained wt HeLa and Drp1 KO HeLa cells with the fluorescently labeled bivD7_AF647_, confirming its functionality as an imaging probe (**Fig. S9**). Additionally, we stained Drp1 in a cell line expressing monomeric enhanced GFP (mEGFP)-tagged Drp1 (mEGFP-Drp1) at endogenous expression levels using the bivD7_AF647_. Confocal imaging and colocalization analysis revealed a reasonable overlap of the mEGFP-Drp1 and the bivD7_AF647_ signal with a Pearsońs correlation coefficient of 0.73 ± 0.07 (**Fig. 5A-C**). Next, we imaged Drp1 stained with bivD7_AF647_ in mEGFP-Drp1 U2OS cells using STORM (**Fig. 5D**). The reconstituted bivD7_AF647_ signal overlaid with the epifluorescence signal of mEGFP-Drp1 acquired in the same cell before STORM imaging as expected. These results demonstrate the applicability of the bivD7_AF647_ as an imaging probe with enough brightness and specificity to label Drp1 at endogenous expression levels. Finally, we imaged bivD7_AF647_-stained Drp1 in wt U2OS cells by STORM. Here, we detected localizations of both diffuse cytosolic Drp1 as well as mitochondrial Drp1 complexes (**Fig. 5E,F**). Utilizing the bivD7_AF647_ for STORM imaging allowed us to resolve macromolecular assemblies of Drp1 at mitochondria demonstrating its usability as a specific Drp1 probe for SRM imaging with minimal linkage error (**Fig. 5G**).

**Figure 5:**
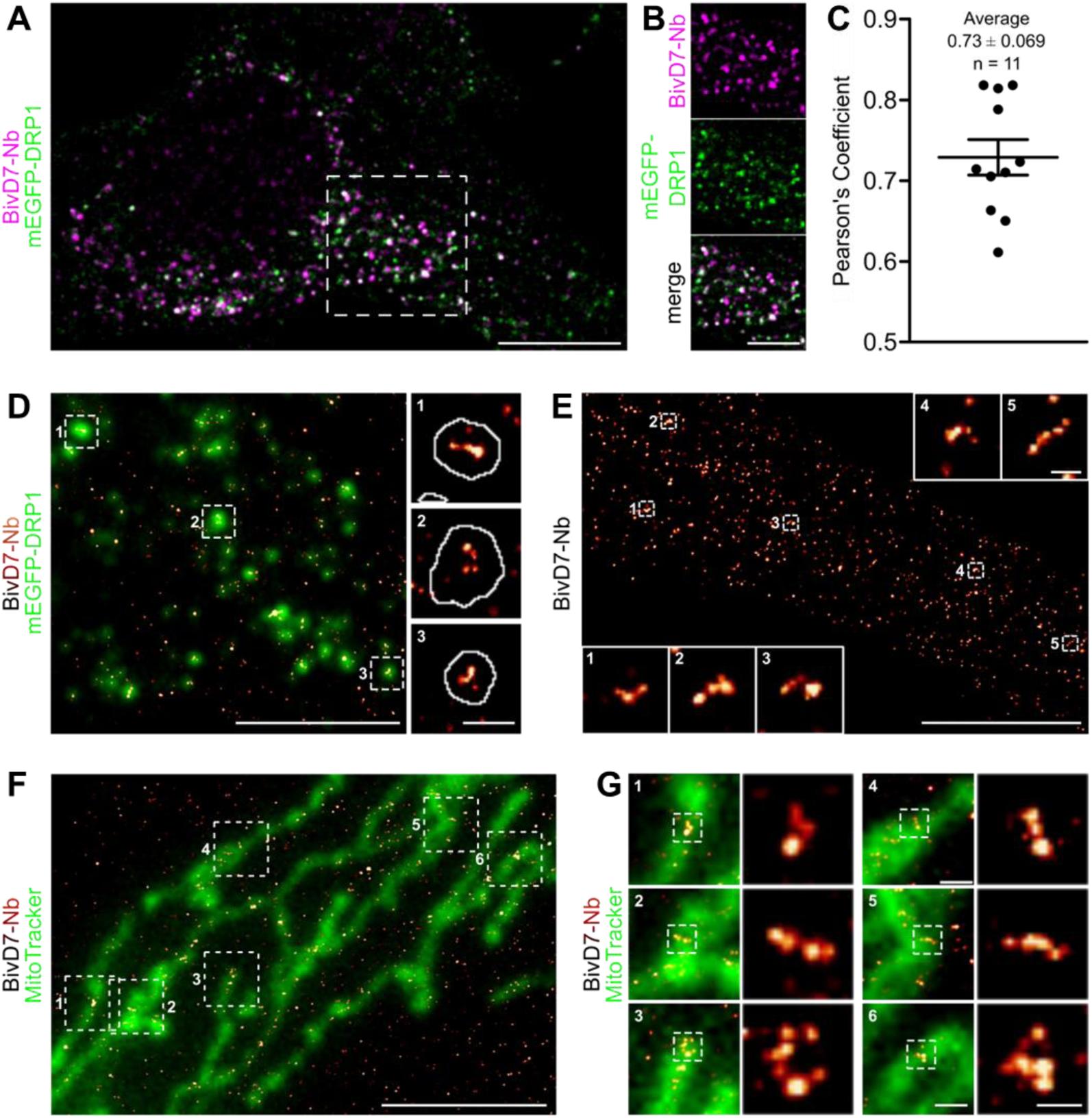
Super-resolution imaging of Drp1 in U2OS cells. (**A**) Representative confocal fluorescence microscopy image of U2OS mEGFP-DRP1 cells (mEGFP-DRP1 signal shown in green) stained with bivD7_AF647_ (magenta). (**B**) Zoomed images correspond to crop regions as indicated in (A). Scale bar 10 µm, crop 5 µm. (**C**) Pearson’s correlation coefficient of mEGFP and bivD7_AF647_ fluorescence emission signals calculated from background-corrected images shown in (**A**). Data are representative for n = 3 independent experiments with n = 11 cells. (**D**) Reconstructed super-resolution image of DRP1 stained with bivD7_AF647_ (orange-hot) overlayed with the epifluorescence signal of mEGFP-Drp1 (green). Zoomed images (right) correspond to cropped regions as indicated. White line in the zoomed regions correspond to the outline of the thresholded mEGFP-DRP1 signal. Images are representative for n = 3 independent experiments. Scale bar 5 µm, zoom 500 nm. (**E**) Reconstructed super-resolution image of DRP1 labelled with bivD7_AF647_ in wt U2OS cells. Insets correspond to zoomed regions as indicated. Scale bar 5 µm, insets 200 nm. (**F**) Reconstructed super-resolution image of Drp1 labelled with bivD7_AF647_ (orange-hot) overlayed with an epifluorescence microscopy image of mitochondria stained with MitoTracker (green) in wt U2OS cells. Scale bar 5 µm. (**G**) Zoomed areas of the image shown in (**F**) (left images) with further zoom on Drp1 assemblies (right images) as indicated. Scale bar 500 nm, zoomed images 200 nm. (**D-G**) Data are representative of at least n = 3 independent experiments.

### Drp1-Cbs visualize the dynamics of endogenous Drp1 in live cells

To visualize endogenous Drp1 in a “tag-free” approach in living cells, we generated Cb expression constructs by genetic fusion of D7 and D63 with TagRFP via a flexible Gly-Ser linker (**Fig. 6A**). First we tested the intracellular binding properties of the newly generated Drp1 Cbs by intracellular immunoprecipitation (IC-IP) (Maier et al., 2015; Traenkle et al., 2015). To this end, we transiently transfected HEK293 cells with the Drp1 Cbs or an unrelated Cb (Pep Cb, (Traenkle et al., 2020)) as a negative control (NC), precipitated the Cbs with a TagRFP nanotrap, and analysed the bound fraction for endogenous Drp1. Immunoblot analysis showed enrichment of endogenous Drp1 along with the precipitated Drp1 Cbs, but not with the negative control (**Fig. 6A**), from which we concluded that both Drp1 Cbs retain their binding properties upon intracellular expression. When using Cbs as intracellular probes, it is important to be aware that the fluorescent emission signal originating from the unbound Cbs may mask the signals of the antigen-bound Cbs. This is particularly a problem for the detection of Drp1, as Drp1 is predominantly localized in the cytosol and unbound cytosolic Cbs result in a high background. This limits the specific detection of Drp1 and e.g. its recruitment to mitochondrial fission sites. To reduce the amount of unbound Cbs and improve the signal-to-noise ratio, we further modified our Drp1 Cbs and generated turnover-accelerated versions that are subjected to faster proteasomal degradation if unbound as previously described (Keller et al., 2018). We tested the applicability of this approach for Drp1 Cbs by expressing both standard and turnover-accelerated D7 and D63 Cbs in wt HeLa and Drp1 KO HeLa cells. Quantitative imaging of living cells using GFP localized in the nucleus for transfection control and cell segmentation revealed significantly lower average cellular intensities and thus a lower cytosolic background originating from the unbound Cb fraction in the case of turnover-accelerated Cbs compared to their unmodified counterpart (**Fig. 6B, C**). Moreover, in Drp1 KO HeLa cells the signal of turnover-accelerated Cbs was further reduced due to the lack of antigen (**Fig. 6B, C**). This decrease indicates that the modification significantly reduces the amount of unbound Cbs and increases the signal-to-noise ratio of these intracellular imaging probes. Finally, we analysed the potency of turnover-accelerated Cbs to visualize the dynamic relocalization of Drp1 from the cytosol to the mitochondria after chemical induction of mitochondrial fission. To induce fission and, thus, recruitment of Drp1 to mitochondria, we treated U2OS cells transiently expressing unmodified or turnover-accelerated Cbs with carbonyl cyanide m-chlorophenylhydrazone (CCCP), an uncoupler of mitochondrial oxidative phosphorylation. Time-lapse imaging of cells expressing the unmodified Cbs showed rather diffuse Cb signals that did not change over time. In contrast, in cells expressing the turnover-accelerated Cbs, an increasing number of spots became visible within the observation period (**Fig. 7**). Based on the colocalization with the Mito Tracker signal, these characteristic spots strongly suggest an accumulation of Drp1 at mitochondrial fission sites. Notably, we did not observe such patterns in Drp1 KO HeLa cells, which further underlines that the detected spots are most likely Drp1 complexes bound by the Drp1 Cbs (**Fig. S10**). In summary, these findings indicated that turnover-accelerated Drp1-specific Cbs are suitable for monitoring and visualizing the dynamic localization of Drp1 at the MOM after induction of mitochondrial fission in living cells.

**Figure 6:**
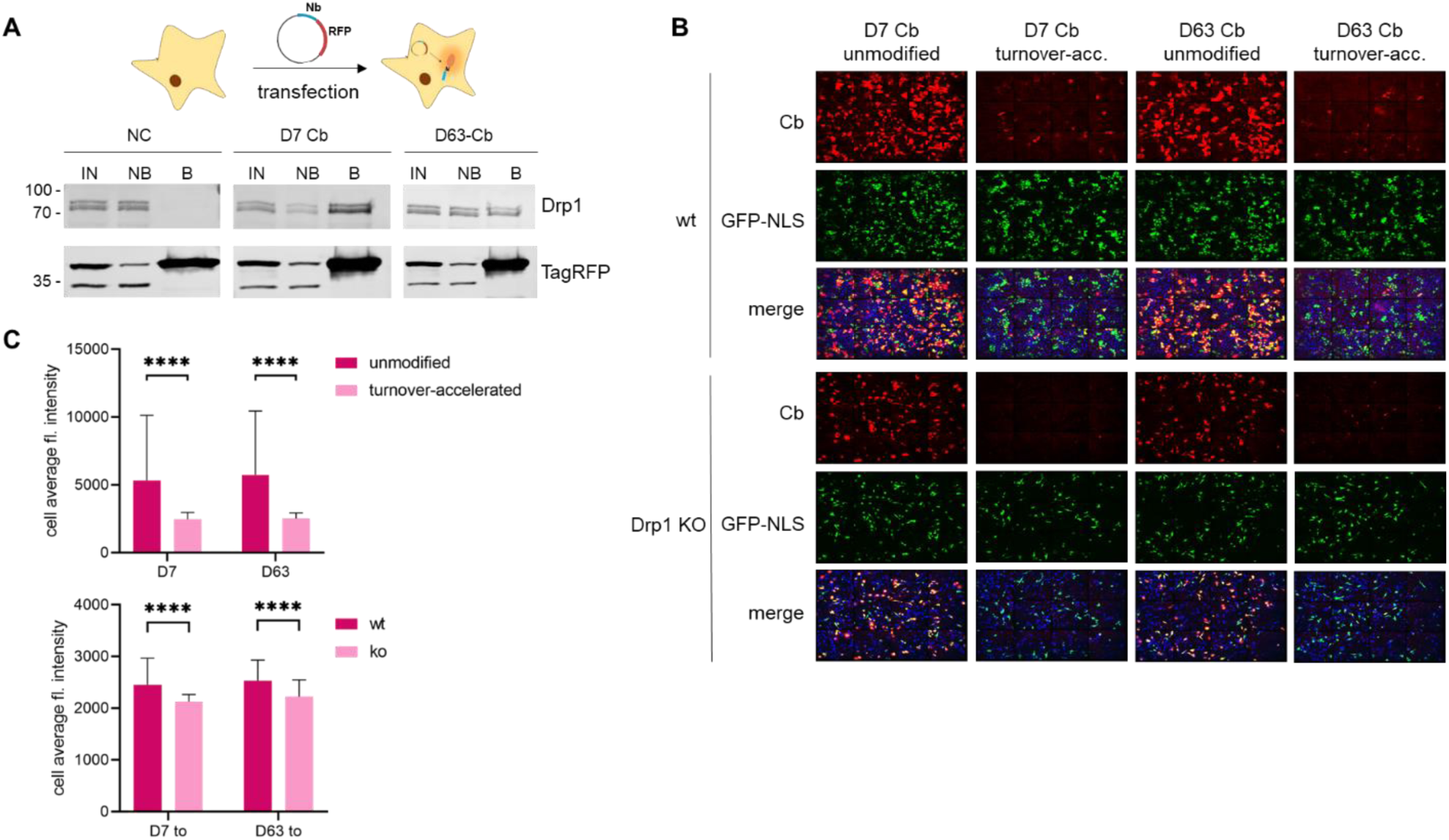
D7-Cb and D63-Cb recognize their antigen and are stabilized in the presence of Drp1 in living cells. (**A**) Intracellular immunoprecipitation (IC-IP) of Drp1. Soluble protein fractions of HEK293 cells either left nontransfected (NC) or expressing indicated chromobodies (D7-CB, D63-CB) were subjected to immunoprecipitation with the RFP-Trap. Input (IN), non-bound (NB) and bound fractions (B) were analysed by immunoblotting with an anti-Drp1 antibody (upper panel) and an anti-TagRFP antibody (lower panel). Molecular weights in kDa on the left. (**B**) Representative fluorescence images of living wt HeLa or Drp1 KO HeLa cells transiently expressing either the unmodified or the turnover-accelerated Cb constructs. As transfection control, a GFP-NLS encoding construct was co-transfected. Nuclei were stained with Hoechst33258. (**C**) Bar chart representing the cell average fluorescence intensity of Cb signal (red channel) quantified from (B). For the comparison of unmodified and turnover-accelerated Cbs, significance was calculated using Kruskal Wallis test with Dunn’s multiple comparisons test (upper panel). For the comparison of the turnover-accelerated Cb intensities in wt HeLa cells and Drp1 KO cells, the significance was calculated using two-way ANOVA with Sidak’s multiple comparisons (lower panel). Data is presented as mean ±SD; n.s. p > 0.05, **** p < 0.0001; n>130 cells each.

**Fig. 7.**
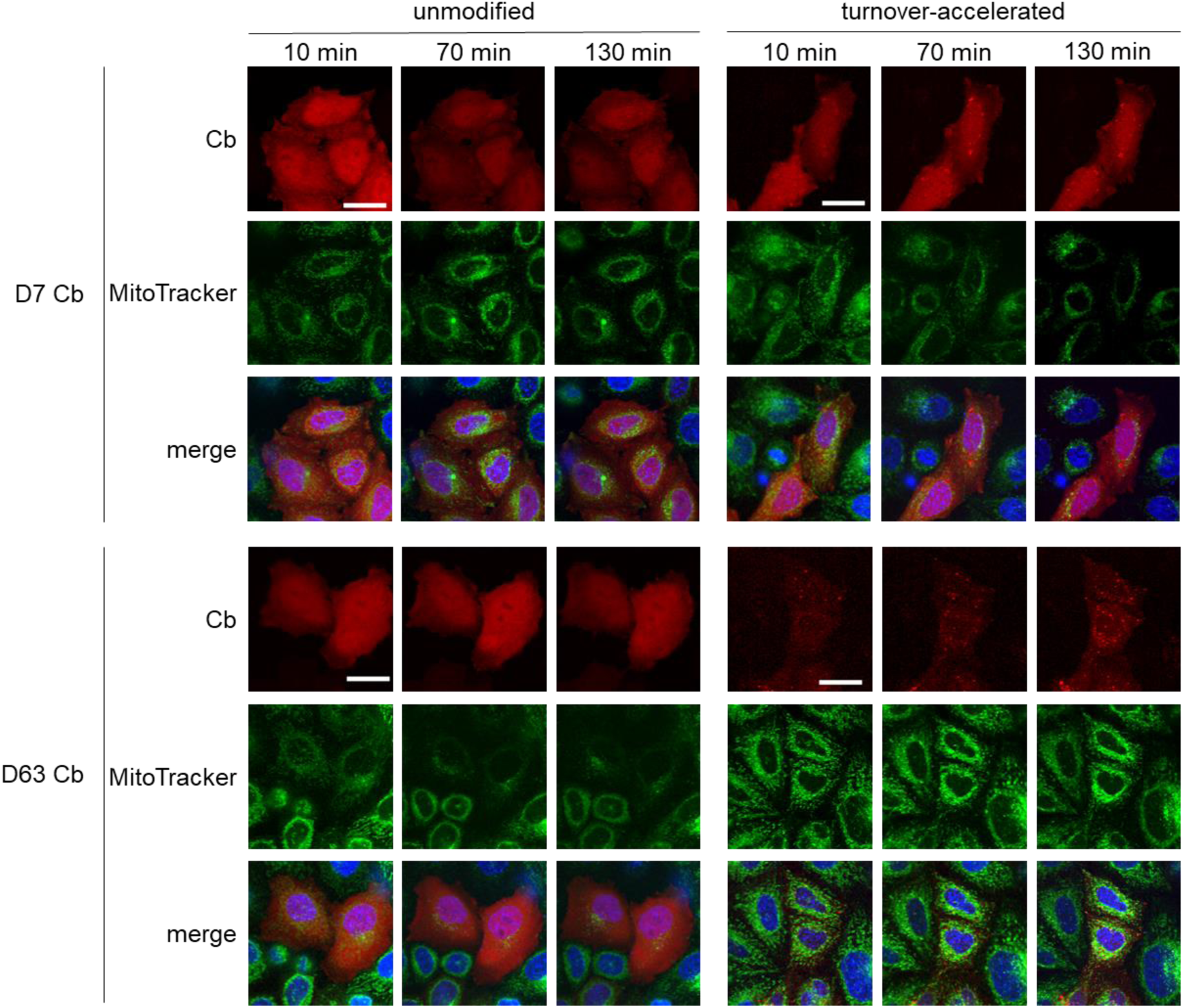
Turnover-accelerated Drp1-Cb monitor relocalization of Drp1 in living cells. Fluorescent time-lapse microscopy of wt HeLa cells transiently expressing unmodified or turnover-accelerated Cbs (red). 24 h post transfection, cells were stained with Hoechst33258 (blue) and MitoTracker green (green) and subsequently treated with 20 µM CCCP. Representative confocal images were taken after 10 min, 70 min, and 130 min of treatment. For each Cb, brightness and contrast were individually set and kept for all time points. Scale bar 25 µm.

## Discussion

There is increasing evidence of dysregulated Drp1 in the context of a variety of diseases, including neurodegeneration such as Parkinson’s (Han et al., 2020), cardiovascular diseases (Kim et al., 2015; Sharp et al., 2014; Zunino et al., 2007) and cancer (Kashatus et al., 2015; Rehman et al., 2012; Serasinghe et al., 2015). However, the exact molecular mechanisms of Drp1 function and its dynamic regulation remain to be elucidated. Therefore, the development of advanced research tools to study this key player in mitochondrial fission is urgently needed (Giacomello et al., 2020; Montecinos-Franjola et al., 2020; Tong et al., 2020; Zerihun et al., 2023).

Here, we present for the first time, to the best of our knowledge, an Nb-based toolset to study endogenous Drp1 in different biochemical and cell biological research settings. We identified two stable and well-producible Nbs, D7 and D63, which bind distinct domains of Drp1 with high affinity. Notably, binding of D63 to the C-terminal GED of Drp1 induces an increased GTPase activity *in vitro*. It can be speculated that this is due to a stabilization of the intramolecular interaction between the GED and the N-terminal GTP-binding domain - a conformation known to be important for the enzymatic function of Drp1 (Frohlich et al., 2013; Zhu et al., 2004). By targeted functionalization and engineering, we have successfully applied the identified Nbs as i) affinity capture tools (Drp1-nanotraps), ii) labelling probes for fluorescence microscopy including SRM, and iii) Cbs for visualization of Drp1 in living cells. For affinity capture from whole cell lysates, we demonstrated that both Drp1-nanotraps efficiently precipitated Drp1. Notably, immunoblot analysis with a monoclonal Drp1 antibody showed captured Drp1 in the form of several bands. These findings are comparable to previous reports also showing multiple bands for Drp1 (Loson et al., 2013; Xie et al., 2020; Yu et al., 2021; Yu et al., 2019) which can be attributed to the size differences between the Drp1 isoforms (Chen et al., 2000; Itoh et al., 2018). Applying the nanotraps to precipitate Drp1 from other cell types or even tissues, might result in the identification of a different Drp1 pattern reflecting the expression of different isoforms. With in-depth MS analysis of the proteome of both Drp1-nanotraps we confirmed the effective and specific enrichment of Drp1 with a high sequence coverage based on the identified peptides. It is noteworthy that the MS analysis also revealed the enrichment of MOM- and peroxisome-associated factors. Interestingly, the latter were found exclusively for the D63-nanotrap, but not for the D7-nanotrap, which could be explained by steric and allosteric effects caused by the binding of the Nbs at the different domains of Drp1. Overall, both Drp1-nanotraps precipitated endogenous Drp1 substantially better compared to the conventional monoclonal Drp1 antibody. Thus, we conceive that both Nb-based affinity matrices enable a fast and efficient investigation of the Drp1-interatome under different physiological conditions and/or in different tissues.

Despite their high affinity for recombinant Drp1, both Nbs initially failed to stain endogenous Drp1 by immunofluorescence labelling. This might be due to a lower accessibility of the epitope or a higher off-rate of the Nbs for Drp1 in its native environment. To improve their binding properties by increasing avidity, we have generated bivalent Nbs, a format that has previously significantly improved antigen recognition for other Nbs (Fagbadebo et al., 2022; Virant et al., 2018). Notably, we found that bivD7 is able to specifically stain endogenous Drp1 in fixed cells. Most importantly, by site-directed fluorophore conjugation we generated a probe with a substantially smaller linkage error compared to conventional antibodies, applicable for super-resolution STORM imaging of Drp1. Commonly applied labelling strategies for SRM include direct (labelled primary antibody) and indirect (labelled secondary antibody) immunofluorescence techniques or genetically encoded fluorescent protein fusions. The linkage error, which describes the distance between the biological sample and the detectable moiety, is determined by the size of the structure used to label the protein of interest (e.g. the antibody) and the random distribution of fluorescence emitters on this structure. This can induce a fluorescence displacement of up to ∼30 nm causing mislocalization of the target structure coordinates (Carrington et al., 2019; Fruh et al., 2021). As demonstrated, the bivD7 conjugated to AlexaFluor647 enables for the first time super-resolved imaging of endogenous Drp1 complexes located in the cytoplasm and on mitochondria.

Recently, it was reported that labelling of Drp1 with genetically-fused tags including fluorescent proteins such as GFP alters its oligomerization state and consequently affects its function (Montecinos-Franjola et al., 2020). Therefore, a tag-free approach is desirable to minimize artefacts when studying Drp1 in living cells. In this context, transiently binding Cbs are known as versatile probes to visualize endogenous antigens in living cells with high spatial and temporal resolution (Burgess et al., 2012; Maier et al., 2015; Panza et al., 2015; Rothbauer et al., 2006; Wegner et al., 2017). Here, we reformatted D7 and D63 into intracellularly functional Cbs and monitored the dynamics of endogenous Drp1 by fluorescent time-lapse imaging of living cells. Considering that Drp1 is mainly diffusely localized in the cytoplasm, we had to face the challenge of distinguishing between the signals originating from Drp1-binding Cbs and unbound Cbs, a problem that is constantly arising when using genetically encoded imaging probes expressed from a constitutive promoter (Wagner and Rothbauer, 2020). Several approaches have been developed to reduce the levels of unbound intrabodies and Cbs (Gross et al., 2013; Keller et al., 2018; Sibler et al., 2005; Tang et al., 2016). Here, we introduced an N-terminal arginine residue, which was previously reported to convey a fast proteasomal degradation of unbound Cbs in the cytoplasm via the N-end rule (Keller et al., 2018; Varshavsky, 2005). The improved signal-to-noise ratio of these turnover-accelerated Drp1 Cbs now enables for the first time the visualization of the drug-induced relocalization of endogenous cytoplasmic Drp1 towards mitochondrial fission sites in living cells. In addition to imaging, the Nbs can be further adapted as building blocks of e.g. Drp1 modulating functional intrabodies for extended research applications. As an example, we are developing Drp1 Nbs as inducible degrons to mediate the targeted degradation of Drp1 in living cells. In summary, currently shown as a proof-of-principle study, we are convinced that the herein-presented Nbs open up new opportunities for comprehensive and detailed studies of Drp1 and its pathophysiological role as a key regulator of mitochondrial fission in advanced experimental settings.

## Material and Methods

### Nanobody library construction

Alpaca immunization with recombinant Drp1 as well as Nb library generation was performed as described before (Rothbauer et al., 2006). Animal immunization was authorized by the government of Upper Bavaria (Permit number 55.2-1-54-2532.0-80-14). Briefly, an alpaca (*Vicugna pacos*) was immunized with recombinant human Drp1 (hDrp1). An initial priming dose of 0.48 mg Drp1 was followed by booster injections of 0.24 mg each in the 3^rd^, 4^th^, 7^th^, and 12^th^ week. Serum samples of 20 ml were taken in the 9^th^ week of immunization for analysis of seroconversion by enzyme-linked immunosorbent assay (ELISA). 13 weeks after the first dose, 100 ml blood were drawn, and the lymphocytes isolated by Ficoll gradient centrifugation using the Lymphocyte Separation Medium (PAA Laboratories GmbH). Total RNA was isolated using TRIzol (Life Technologies) followed by reverse transcription of mRNA into cDNA using a First-Strand cDNA Synthesis Kit (GE Healthcare). To isolate the Nb repertoire, 3 nested PCR reactions were performed with the following primers: (i) CALL001 and CALL002, (ii) forward primer set FR1-1, FR1-2, FR1-3, FR1-4 and reverse primer CALL002, and (iii) forward primer FR1-ext1 and FR1-ext2 and reverse primer set FR4-1, FR4-2, FR4-3, FR4-4, FR4-5 and FR4-6 to introduce SfiI and NotI restriction sites. The sequences for all primers (synthesized by Integrated DNA Technologies (Leuven, Belgium)) used in this study are listed in **Table S4**. Then, the Nb library was subcloned into the pHEN4 phagemid vector (Arbabi Ghahroudi et al., 1997) using the Sfil/NotI restriction sites.

### Nanobody screening

To select for Drp1-specific Nbs, *E. coli* TG1 cells containing the Drp1-Nb library in pHEN4 were first infected with the M13K07 helper phage. Next, the hereby generated Nb-presenting phages were isolated from culture supernatant by PEG precipitation and 1×10^11^ phages were used for subsequent panning. In each selection round, extensive blocking of both recombinant Drp1 and phages was carried out with 5 % milk or BSA in PBST (PBS, 0.05 % Tween 20, pH 7.4) (Pardon et al., 2014). To deplete non-specific binders, phages were first added to Nunc Immuno MaxiSorp tubes (Thermo Scientific) coated with 10 µg/ml GFP before transferring them to tubes coated with 10 µg/ml Drp1 or 10 µg/ml GFP as non-related antigen for 2 hours at room temperature. Next, unbound phages were removed by washing with stringency increasing each round. Bound phages were eluted with 100 mM triethylamine (pH 12.0) and immediately neutralized with 1 M Tris/HCl pH 7.4. After each round of panning, log phase *E. coli* TG1 cells were infected with the eluted phages and grown on selection plates to rescue enriched phages. Drp1-specific enrichment was monitored by determining the colony-forming units (CFUs) after each round. After two rounds of panning, 260 individual clones were randomly selected and screened by phage ELISA using immobilized Drp1 or GFP as control. Horseradish peroxidase-labeled anti-M13 monoclonal antibody (GE Healthcare) and Thermo Scientific 1-Step Ultra TMB solution were used for detection of bound phages. The reaction was stopped with 100 µL of 1 M H_2_SO_4_ and the signal was detected with the Pherastar plate reader at 450 nm. Phage ELISA-positive clones were defined by a 2-fold signal above the control.

### Expression plasmids

For bacterial Nb expression, Nb encoding cDNA were cloned into the pHEN6C vector as previously described (Rothbauer et al., 2008). For mammalian expression of bivalent Nbs, cDNAs comprising the two coding sequences of the respective Nbs connected by a flexible Gly-Ser linker [(G_4_S)_4_] and a C-terminal sortase tag (L-P-E-T-G) followed by a His_6_-tag were generated and inserted into pCDNA3.4 expression vector variant comprising N-terminal signal peptide for secretion (Wagner et al., 2021) as following: For bivD7, each Nb sequence was amplified separately by PCR using bivD7-1 and bivD7-2 or bivD7-3 and downEcoRI-rev primer pairs. Next, both Nbs were fused by overlap-extension PCR using the primers bivNshort_for and downEcoRI-rev. For bivD63 cloning, the primer pairs bivD63forN and bivNtermGS-rev as well as bivD63forC and downEcoRI-rev were used followed by bivD63_fusion and downEcoRI-rev for overlap-extension PCR. Finally, the resulting sequence was inserted into pCDNA3.4 via restriction digestion with Esp3I and EcoRI. For unmodified Cbs, Nb sequences were cloned into pTagRFP by BglII and HindIII restriction digest as previously described (Panza et al., 2015). For turnover-accelerated Cbs, the Nb sequence was inserted via PstI and BspEI into the previously described pEGFP-Ubi-R-3xGS-tagRFP plasmid (Keller et al., 2018). The coding sequence of Drp1 was amplified from pAcGFP-Drp1 plasmid (Jenner et al., 2022) with Drp1_GTPaseFor and Drp1_C_term_rev and cloned into pEGFP-N1 backbone. For deletion constructs for domain mapping all sequences were amplified from AcGFP-Drp1 and cloned into pEGFP-N1 with Drp1-GTPaseFor forward primer and the respective reverse primer: Drp1-VD-rev (GTPase-MD-VD), Drp1-MD-rev (GTPase-MD), Drp1-GTPase-rev (GTPase). The correct sequences of all constructs were confirmed by Sanger sequencing. All expression constructs used in this study are listed in **Table S5**.

### Protein expression and purification

Monovalent Drp1 Nbs were expressed and purified as previously described (Burgstaller et al., 2022). Bivalent Drp1 Nbs were produced using the ExpiCHO system (Thermo Fischer Scientific) according to the manufacturer’s instructions (Fagbadebo et al., 2022). Purity of produced Nbs was assessed using SDS-PAGE. For this, protein samples were denatured (10 min, 95 °C) with 2x SDS-sample buffer (60 mM Tris/HCl, pH 6.8; 2 % (w/v) SDS; 5 % (v/v) 2-mercaptoethanol; 10 % (v/v) glycerol, 0.02 % bromphenol blue) prior to loading onto the gel. To visualize the total protein, gels were stained with InstantBlue Coomassie (Abcam) and protein concentrations were determined using Bradford and spectrophotometer measurements. hDrp1 was purified as described in (Jenner et al., 2022). Briefly, hDRP1 (isoform 1) was expressed from a pTYB2 vector in *E. coli* BL21-CodonPlus (DE3)-RIPL. Bacteria were grown at 37 °C in LB medium containing 100 µg/ml ampicillin and 35 µg/ml chloramphenicol to an OD_600 nm_ of 0.6. Protein expression was induced using 1 mM isopropyl 1-thio-β-d-galactopyranoside (IPTG) at 14 °C for 18 hours. Cell pellets were resuspended in 20 mM HEPES/KOH pH 7.4, 500 mM NaCl and 1 mM MgCl_2_ containing 1 mM PMSF, 1 µg/ml DNAse I (Roche Diagnostics), and protease inhibitor (Complete; EDTA-free Protease Inhibitor Mixture; Roche Applied Science), homogenized by mechanical rupture and cell debris was removed by centrifugation. The supernatant was purified using chitin resin affinity purification (New England Biolabs, Inc.). hDrp1 was eluted by cleavage from the affinity resin using 30 mM DTT at pH 8 for 48 hours at 4 °C and further purified using anion exchange chromatography (HiTrap Q FF, GE Healthcare) at pH 8. Pure elution fractions were pooled and dialyzed against 20 mM HEPES/KOH pH 7.4, 500 mM NaCl and 1 mM MgCl_2_. Purity was assessed using SDS-PAGE as described and protein concentration was determined using Bradford and spectrophotometer measurements. Purified protein was stored with 20 % (v/v) glycerol.

### Affinity Measurements by biolayer interferometry

Binding kinetics analysis of Drp1 Nbs was performed using the Octet RED96e system (Sartorius) according to the manufacturer’s recommendations. In brief, streptavidin-coated biosensor tips (SA, Sartorius) were incubated for 40 s with 1.7-10 µg/ml biotinylated Drp1-Nbs diluted in Octet buffer (HEPES, 0.1 % BSA). In the association step, a 4-step 2-fold dilution series of Drp1 starting from 125 nM (D7), 62.5 nM (D63) or 120 nM (bivD7 and bivD63) were applied for 300 s followed by dissociation in Octet buffer for 720 s. Each concentration was normalized to a reference applying Octet buffer only for association. Data were analysed with the Octet Data Analysis HT 12.0 software using the 1:1 ligand model and global fitting.

### Drp1 GTPase activity measurements

GTPase activities for Drp1 were measured using a continous, regenerative assay described by (Ingerman and Nunnari, 2005). In brief, Drp1 (100 nM final) was incubated with 0, 10, 100 or 1000 nM GFP-Nb, D7, or D63. Upon addition of 1 mM DTT, 1 mM PEPK, 600 µM NADH, ≥20 U/ml PK/LDH (all Sigma Aldrich) and 1 mM GTP, GDP (both Jena Bioscience), or no nucleotide, the reactions were started with the final buffer containing 20 mM Hepes pH 7.4, 150 mM KCl, 2 mM MgCl_2_. Upon thermal equilibration of the microtiter plates, NADH absorbances were measured for 120 min using a Safire Tecan-F129013 microplate reader (Tecan, Austria) operating at 340 nm and 37 °C. Using BSA without nucleotide or GDP for rapid NADH depletion, absorbances were converted into NADH concentrations. GTPase rates were calculated using linear fits between 30 and 120 min of the traces.

### Cell culture and transfections

U2OS and HEK293 cell lines were obtained from ATCC (CRL3216, HTB-96); the HeLa Kyoto cell line (Cellosaurus no. CVCL_1922) was acquired from S. Narumiya (Kyoto University, Japan). Drp1 KO HeLa cells were generated in and provided by the lab of Thomas Langer (Max Plack Institute, Cologne, Germany). For mycoplasma testing, the mycoplasma kit Venor GeM Classic (Minerva Biolabs) and Taq polymerase (Minerva Biolabs) were applied. No additional authentication was performed as this study does not contain any cell-line specific experiments. Culturing of cell lines was carried out by standard protocols. In brief, cells were grown in DMEM (high glucose, pyruvate, GlutaMax (Thermo Fischer Scientific) with 10 % (v/v) fetal calf serum (Thermo Fischer Scientific) and 1 % (v/v) penicillin/streptomycin (Thermo Fischer Scientific). For routine passaging of cells, 0.05 % trypsin-EDTA (Thermo Fischer Scientific) was used. Cells were cultivated at 37 °C and 5 % CO_2_ in a humidified incubator. U2OS cells were transiently transfected with Lipotectamine2000 (Thermo Fisher Scientific) according to the manufacturer’s recommendations. HEK293 and HeLa cells were transfected with Polyethyleneimine (PEI, Sigma Aldrich) as previously described (Braun et al., 2016).

### Nanobody immobilization on NHS-sepharose matrix

1.2 mg Drp1-Nbs per 1 ml NHS-Sepharose (Cytiva) were immobilized according to the manufacturer’s instructions.

### Sortase labelling of nanobodies

For sortase coupling, 50 μM Nb, 250 μM sortase peptide (H-Gly-Gly-Gly-propyl-azide, synthesized by AG Maurer, University of Tübingen) dissolved in sortase buffer (50 mM Tris, pH 7.5, and 150 mM NaCl) and 10 μM sortase were mixed in coupling buffer (sortase buffer with 10 mM CaCl_2_) and incubated for 4 h at 4 °C. Uncoupled Nb and Sortase were removed by IMAC, and the excess peptide was depleted via Amicon Ultra Centrifugal Filters (Merck Millipore). To label the azide-coupled nanobody with a fluorophore, SPACC (strain-promoted azide-alkyne cycloaddition) click chemistry reaction was applied. For this, 5-fold molar excess of DBCO-AF647 (Jena Bioscience) was added to the azide-coupled Nb followed by incubation for 2 hours at room temperature and subsequently purified by dialysis and hydrophobic interaction chromatography (HIC).

### Mammalian cell lysis and protein extraction

For intracellular immunoprecipitation (IC-IP), 2.5 – 3 x 10^6^ HEK293 cells were seeded in a 100 mm culture dish (P100). The next day, cells were transfected with the chromobody-encoding plasmid. After 24 hours, cells were harvested at 90 % confluency. This workflow was also followed for the domain mapping constructs. For immunoprecipitations (IPs) using nanotraps, T175 flasks or P100 dishes of HEK293, U2OS or HeLa cells were harvested at ∼90 % confluency. HEK293 cells were detached by light pipetting and U2OS or HeLa cells by trypsination. Subsequently, the cells were washed with 1 ml PBS, and the pellets were stored at −80 °C until lysis. For lysis, T175 flask pellets were resuspended in 600 µL, P100 pellets in 200 µL lysis buffer (50 mM Tris HCl, pH7.5; 150 mM NaCl; 1 mM EDTA, pH 8; 0.5 % TritonX-100; 1 µg/µL DNaseI; 2.5 mM MgCl_2_; 2 mM PMSF;1x Protease inhibitor Mix M (Serva)). The lysate was passed through 20G, 23G and 27G needle gauges multiple times with intermediate vortexing and 10 min incubation steps on ice. Subsequently, the lysate was centrifuged for 10 min at 18000 g at 4 °C. The supernatant was transferred to a fresh tube and diluted 1.5-fold with dilution buffer (50 mM Tris HCl pH 7.5; 150 mM NaCl; 1 mM EDTA, pH 8; 2 mM PMSF).

### Immunoprecipitation

Soluble protein extracts were generated as described above. For analysis of the input (IN), 20 µL of the lysate were mixed with an equal amount of 4x SDS-Sample buffer. For IP, 80 µL slurry of each Nb-trap was equilibrated in IP buffer (50 mM Tris HCl pH 7.5; 150 mM NaCl; 1 mM EDTA, pH 8) and added to equal amounts of lysate. As positive (domain mapping) or negative control (precipitation of endogenous Drp1), 80 µL GFP-Trap slurry (ChromoTek) was used. For IC-IP, 80 µL of RFP-Trap slurry (ChromoTek) was used. Soluble protein extracts were incubated with the Nb-traps overnight at 4 °C on an-end-over end rotor. For positive control (precipitation of endogenous Drp1), 4 µL Drp1-antibody was added to the lysate and after overnight incubation at 4 °C 80 µL equilibrated Protein A/G sepharose slurry were added followed by additional 3-4 hours incubation at 4 °C. Beads were harvested by centrifugation (2000 g, 2 min, 4 °C) and supernatant (non-bound sample) was mixed 1:1 with 4x SDS-sample buffer. The beads were washed 3 times with 500 µL wash buffer (IP buffer with 2 mM PMSF, for MS samples additionally 1x Protease inhibitor mix and 0.05 % Tween20). With the third wash, beads were transferred into a fresh tube. The beads were boiled in 60 µL 2x SDS-sample buffer for 10 min at 95 °C and the supernatant was transferred into a fresh tube (bound sample).

For immunoblotting of IP samples, 1 % of input and non-bound and 33 % of the bound sample were loaded on an SDS-Page and blotted on nitrocellulose membrane as previously described (Fagbadebo et al., 2022). Immunoblots were probed with the following antibodies: anti-Drp1, anti-TagRFP, anti-GAPDH (all antibodies used in this study are listed in **Table S6**). For immunoblot scanning, a Typhoon-Trio laser scanner (GE Healthcare) was used with excitation 633 nm and emission filter 670 nm BP 30 settings.

### Chromobody imaging

U2OS cells were seeded at 8000 cells/well in µclear 96 well plates (Greiner Bio-One). Wt HeLa cells were seeded at 8000 cells/well, Drp1 KO HeLa cells at 11000 cells/well. The next day, cells were transfected with the respective plasmids. 24 hours post transfection, cells were stained with 400 nM MitoTracker green (life technologies) and 4 µg/ml Hoechst33258 for 30 min. Afterwards, medium was replaced with live-cell visualization medium DMEMgfp-2 (Evrogen) supplemented with 10 % FBS, 2 mM L-glutamine with or without 20 µM Carbonyl cyanide m-chlorophenylhydrazone (CCCP). Images were taken 10 min, 70 min and 130 min after compound addition with ImageExpress Micro Confocal High Content screening system (Molecular devices) at 40x magnification. During imaging, cells were kept at 37 °C and 5 % CO_2_. Raw microscopy images were adjusted in brightness and contrast using MetaExpress software (64 bit, 6.2.3.733 or higher, Molecular devices).

### Immunofluorescence in fixed cells

Cells were seeded in µclear 96 well plates (Greiner Bio-One) in the same densities as described for Cb imaging. The next day, cells were transfected with GFP-Drp1 plasmid or left untransfected. 24 hours post transfection, cells were washed twice with PBS and subsequently fixed with 3.7 % paraformaldehyde (PFA) for 10 min at RT. Following three times of washing with PBS, cells were incubated with TBST for 10 min at RT. After additional washes, cells were blocked with 5 % BSA in TBST for 1 hour. Afterwards, the cells were incubated with 100-1000 nM Nb and anti-Drp1 antibody (1:100-1:300) in 5% BSA/TBST overnight at 4°C. Then, unbound Nb and antibody were washed away and the cells incubated with secondary antibody (anti-rabbitAlexaFluor647 (1:1000) and Cy5-conjugated goat anti-alpaca antibody (1:500) for one hour at RT in 2.5 % BSA/TBST. Nuclei staining with 4’, 6-diamidino-2-phenylindole (DAPI, Sigma-Aldrich) was done concurrently. After additional washing with TBST and PBS, images were acquired using an ImageExpress Micro Confocal High Content screening system (Molecular devices) at 40x magnification. Raw microscopy images were adjusted in brightness and contrast using MetaExpress software (64 bit, 6.2.3.733 or higher, Molecular devices).

### Liquid chromatography-MS sample preparation and measurement

Drp1 pulldown using the D7 and D63 nanotraps were compared to the GFP-Trap (negative control) in three replicates. Immunoprecipitation was performed as described above and proteins were subsequently separated by SDS-PAGE (4-12% NuPAGE tris gel (Invitrogen)) for 7 min at 200 V and stained with Coomassie blue. The protein gel pieces were subjected to tryptic digestion as described previously (Shevchenko et al., 2006). Purified peptide samples were measured on a Q Exactive HF mass spectrometer (Thermo Fisher Scientific) online-coupled to an Easy-nLC 1200 UHPLC (Thermo Fisher Scientific). Peptides were separated using a 20-cm-long, 75–μm–inner diameter analytical HPLC column (ID PicoTip fused silica emitter; New Objective, Berks, UK) packed in-house with ReproSil-Pur C18-AQ 1.9-μm silica beads (Dr Maisch GmbH) and eluted using a 60 min segmented gradient from to 10 - 90 % of solvents A (0.1 % formic acid) and B (80 % acetonitrile in 0.1 % formic acid) at a constant flow rate of 200 nl/min. The column temperature was maintained at 40 °C using an integrated column oven. Peptides were ionized by nanospray ionization and a capillary temperature of 275 °C. The mass spectrometer was operated in the positive ion mode. Full MS scans were acquired in a range of 300-1750 m/z at resolution of 60,000. Seven most intense multiple-charged ions were selected for HCD fragmentation with a dynamic exclusion period of 30 s and tandem MS (MS/MS) spectra were acquired at resolution of 15,000.

### Mass spectrometry data analysis

The acquired raw files were processed by MaxQuant software (version 2.0.3.0.) (Cox and Mann, 2008) and searched against the Uniprot *Homo sapiens* database (105.079 entries, downloaded 2022/08/01), *Vicugna pacos* derived nanotraps and commonly observed protein contaminants. For MS and MS/MS, the peptide mass tolerance was set at 4.5 ppm and 20 ppm, respectively. Only two missed cleavages were allowed for the tryptic digestion. Carbamidomethylation (C) was used as fixed modification, while oxidation (M) and acetylation (protein N-term) were defined as variable modifications. False discovery rate was set to 1 % at both peptide and protein level. For label-free quantification a minimum ratio count of two was requested. Intensity-based absolute quantification (iBAQ) was enabled. Downstream analysis of the ‘proteinGroups.txt’ output table was performed in Perseus (version 1.6.15.0). Contaminants, reversed proteins and proteins only identified by site were filtered out. Only proteins that were quantified in two out of three replicates of either the bait or control pulldown were retained in the dataset. Missing values were imputed (width 0.3, down shift 1.8). In order to determine significantly enriched proteins, a multi-volcano analysis (Hawaiian plot) was performed. The *s*_0_ and FDR parameters of the multi-volcano analysis were: for class A (higher confidence, *s*_0_ = 0.1, FDR = 0.01) and class B (lower confidence, *s*_0_ = 0.1, FDR = 0.05). Additional graphical visualization was performed in the R environment (version 4.1.1) and in GraphPad (version 8.0.1.)

### Cell cultivation, seeding and staining for super resolution microscopy

U2OS wild type or U2OS mEGFP-Drp1 cells were cultivated in DMEM (low glucose) supplemented 10 % (v/v) FBS and 1 % (v/v) penicillin/streptomycin (Invitrogen) at 37 °C and 5 % (v/v) CO_2_. For microscopy, the cells were seeded in 35 mm µ-Dish 1.5H glass bottom imaging chambers (for confocal imaging) or on 35 mm 1.5H glass coverslips in in a 6-well plate (for SMLM) at a density of 3 x 10^5^ cells per dish/well and grown for 18 hours. If required, mitochondria were stained using 200 nM MitoTracker Green FM (Invitrogen) for 20 min. Cells were fixed in 4 % (v/v) PFA in DMEM (pre-warmed to 37 °C) for 10 min at room temperature and washed twice with PBS. To quench unreacted fixative, the cells were incubated in 50 mM NH_4_Cl for 15 min followed by permeabilization in 0.25 % (v/v) Triton X-100 for 8 min. The cells were washed with PBS three times for 5 min and incubated with 1 % (w/v) BSA in PBS for 1 hour to block unspecific binding sites. DRP1 was labeled using 250 nM bivD7 coupled to AlexaFluor 647 (Invitrogen) in 1 % (w/v) BSA in PBS over night at 4 °C protected from light. The cells were washed three times with PBS and stored at 4 °C in the dark until imaging.

### Confocal fluorescence microscopy

U2OS mEGFP-Drp1 cells were prepared as described above. Confocal fluorescence microscopy images were acquired on an inverted Infinity Line confocal dual-color STED 775 QUAD laser scanning microscope (Abberior Instruments) equipped with a UPLXAPO 60x/1.42 Oil objective (Olympus). mEGFP was excited at 488 nm and AF647 was excited at 640 nm wavelength. Images were acquired with a pixel size of 65 nm and a pixel dwell-time of 5 µs. The fluorescence emission signal was collected on avalanche photodiode detectors. Raw microscopy images were adjusted in brightness and contrast using Fiji/ImageJ (Schindelin et al., 2012). Colocalization of endogenous mEGFP-Drp1 and bivD7_AF647_ was assessed by calculating the Pearsońs correlation coefficient of the respective emission signals using Just Another Co-localization Plugin (JACoP) (Bolte and Cordelières, 2006) in Fiji/ImageJ (Schindelin et al., 2012). For this, the images were cropped to remove unspecific bivD7 signal outside of the cell and in the nucleus. The experiments are representative for n = 3 individual replicates with n = 11 cells in total.

### Single-molecule localization microscopy

U2OS wild type cells were seeded and stained as described above and mounted on custom-made imaging chambers or single concave depression microscopy slides covered with 100-300 µL imaging buffer (50 mM Tris/HCl pH 8, 10 mM NaCl, 10 % (w/v) glucose, 35 mM cysteamine (MEA), 0.5 mg/ml glucose oxidase (Sigma) and 40 μg/ml catalase (Sigma)) and sealed using Twinsil speed dental glue (Picodent, Germany). The samples were imaged on a home-built wide-field/TIRF microscope or on a SR GSD (3D) inverted DMI6000 B super-resolution wide-field/TIRF microscope (Leica Microsystems) equipped with a 100x, 1.47 N.A. GSD Objective, 405 nm, 488 nm, 532 nm and 642 nm excitation lasers and an Andor iXon Ultra 897 EMCCD camera. Image series were acquired with an exposure time between 30 and 50 ms and an EM gain of 300. AF647 was excited using 640 nm wavelength and the 405 nm activation laser intensity was manually adjusted to keep a constant number of localizations per frame. Typically, 70,000–100,000 frames were recorded. Analysis was performed using the LAS X Softeware (Leica) and ThunderStorm (Ovesny et al., 2014). Images were rendered using a Gaussian with a width according to the localization precision. Image analysis was done using Fiji/ImageJ (Schindelin et al., 2012). Images are representative for n = 6 individual experiments.

### Image analysis and statistics

Image analysis was performed with MetaExpress software (64 bit, 6.2.3.733 or higher, Molecular devices) for n>130 cells per construct. Using the Custom editor (version 2.5.13.3 or higher) or the MetaExpress software, we developed an algorithm to determine the average signal intensity per transfected cell. Based on the assumption that cells usually are transfected with both constructs, the GFP-NLS signal was used as a marker for transfection and cell area. Cell average intensity of the TagRFP signal was determined for all GFP-positive cells. Using the DAPI signal in GFP-NLS negative cells, the TagRFP signal intensity was measured for background determination. The average background signal was determined (n=2125 cells) and subtracted from all measurements. A workflow is shown in **Fig. S11**.

Mean and standard deviation were calculated for each condition. For comparison of unmodified and turnover-accelereated Cbs, significance was calculated using Kruskal Wallis test with Dunn’s multiple comparisons test. For comparison of turnover-accelerated Cbs in wt HeLa and Drp1 KO HeLa cells, significance was calculated using two-way ANOVA with Sidak’s multiple comparisons test. Tests were done using GraphPad Prism software (Version 8 or higher). All graphs were prepared using GraphPad Prism.

## Author contributions

T.F. and U.R. conceived the study and analysed the data. T.F., D.I.F., F.O.F., P.D.K., Y.L. and E.S. perform all cellular and biochemical experiments. S.N. and A.S. immunized the alpaca. A.J. and A.G-S purified Drp1 for immunization and performed and analysed super-resolution microscopy experiments. C.C-R. and B.M. performed and analysed mass spectrometry experiments. T.F. and U.R. wrote the manuscript with the help of all authors.

## Acknowledgements

This research was supported by the German Research Foundation (DFG) through RTG 2364 “MOMbrane” to T.F., A.J.; A.G-S., C.C-.R., F.O.F.,Y.L., E.S., B.M and U.R. The authors also gratefully acknowledge the Ministry of Science, Research and Arts of Baden-Württemberg (V.1.4.-H3-1403-74) for financial support. We thank the CECAD imaging facility for excellent assistance.

## Supplementary Information

### Supplementary figures

**Supplementary Figure 1:**
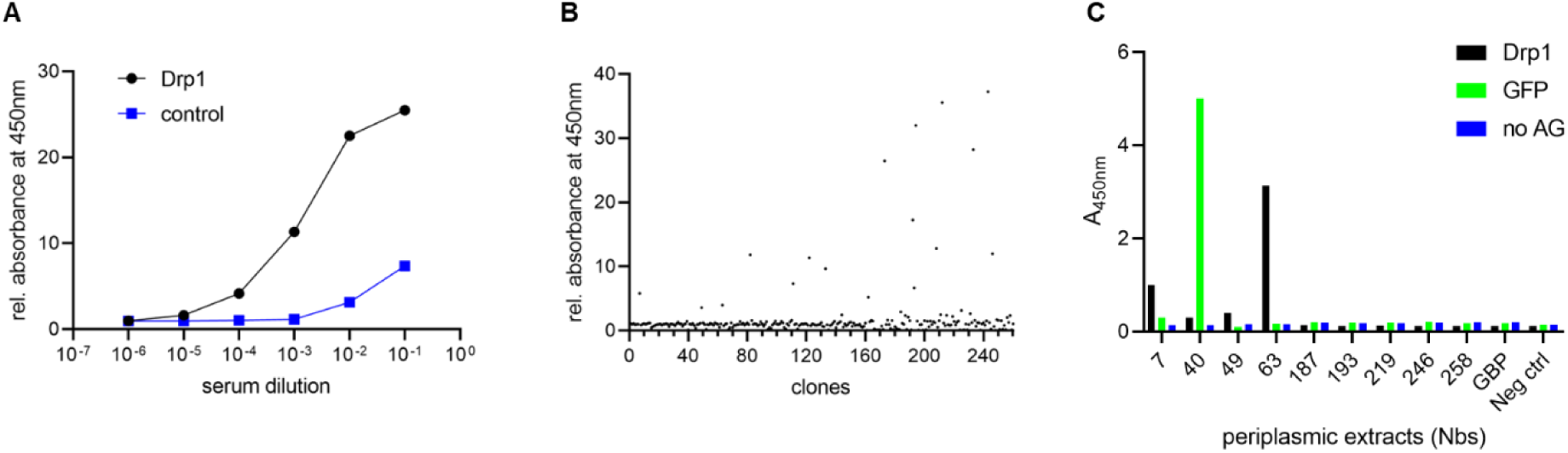
Screening for Drp1-specific nanobodies. (**A**) Analysis of seroconversion upon vaccination with Drp1. A serum sample of the vaccinated alpaca (*Vicugna pacos*) was collected 13 weeks after the first dose. To test for Drp1-specific antibodies, a serum ELISA was performed with the indicated dilutions in plates coated either with Drp1 (black) or bovine serum albumin (BSA, blue) as a negative control. Binding of Drp1-specific antibodies was detected by using an anti-heavy chain antibody conjugated to horseradish peroxidase. (**B**) Result of phage ELISA of 260 selected clones to test for binding of Drp1 immobilized on plates. Signal intensities were normalized to GFP immobilized on plates as negative control. (**C**) Protein ELISA with periplasmic extracts from the eight Nbs identified by phage ELISA.

**Supplementary Figure 2:**
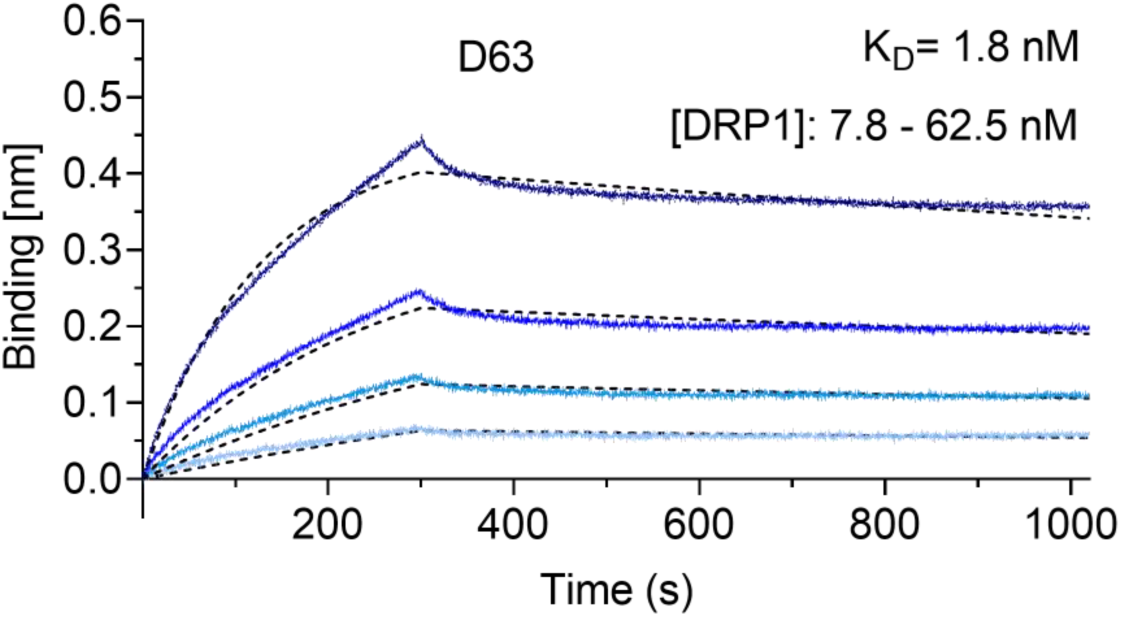
Affinity measurement of D63 Nb. Biolayer interferometry (BLI)-based affinity measurements for D63. Biotinylated D63 was immobilized on streptavidin sensors. Kinetic measurements were performed using four concentrations of purified Drp1 ranging from 7.8 to 62.5 nM (displayed with gradually darker shades of color). The binding affinity (K_D_) was calculated from global 1:1 fits shown as dashed lines.

**Supplementary Figure 3:**
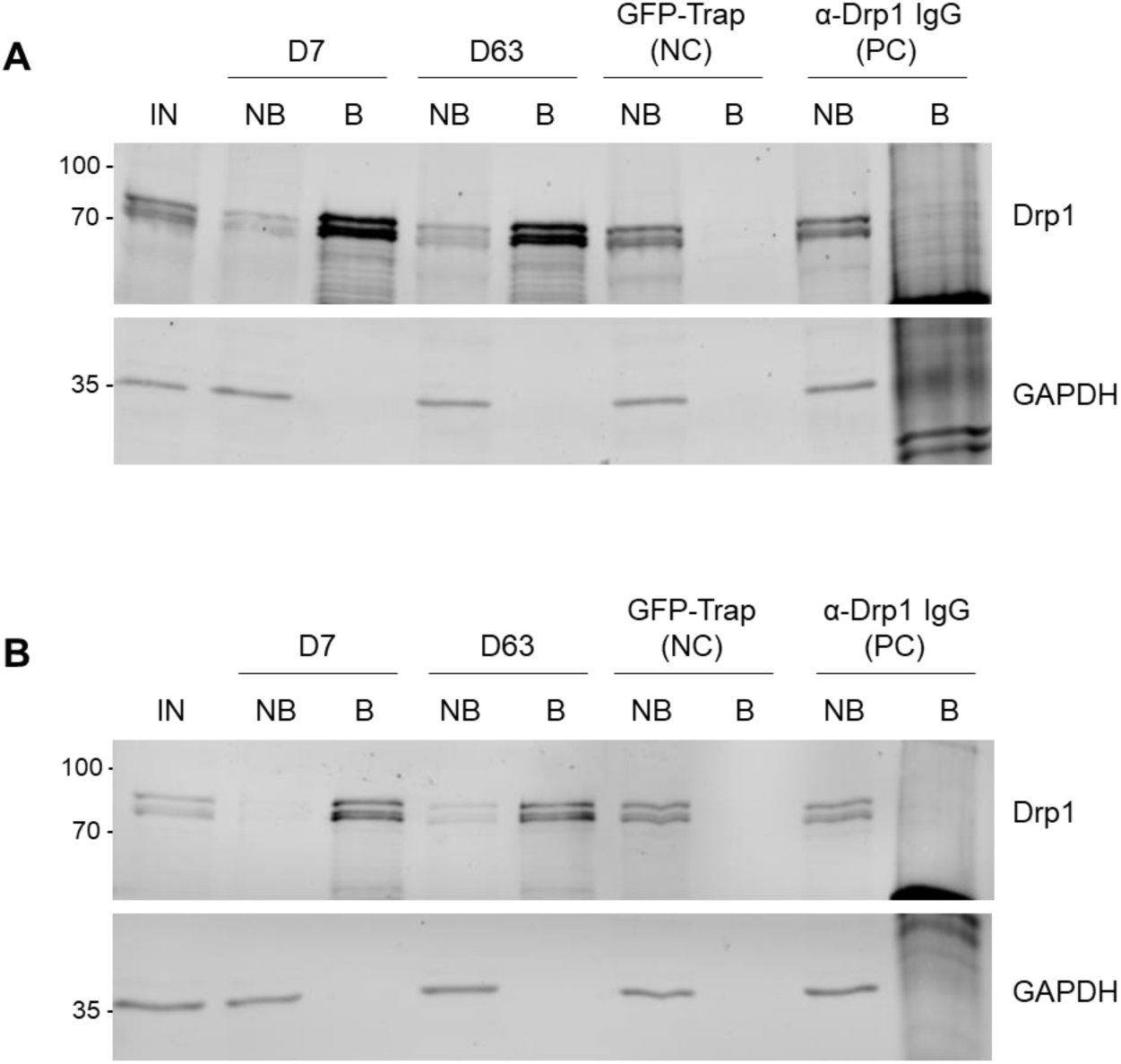
Pulldown of Drp1 from HEK293 and U2OS cells. For immunoprecipitation with nanotraps, the soluble protein fraction of HEK293 cells (**A**) or U2OS cells (**B**) was incubated with equal amounts of the Drp1 nanotraps (D7, D63). As negative control (NC), the GFP-Trap as non-specific nanotrap was used. As positive control, anti-Drp1 IgG was immobilized on protein A/G Sepharose beads (PC). Input (IN, 1 % of total), non-bound (NB, 1 % of total) and bound (B, 33 % of total) fractions were subjected to SDS-PAGE followed by immunoblot analysis using antibodies specific for Drp1 (upper panel) and GAPDH (lower panel). Molecular weights in kDa are indicated on the left.

**Supplementary Figure 4:**
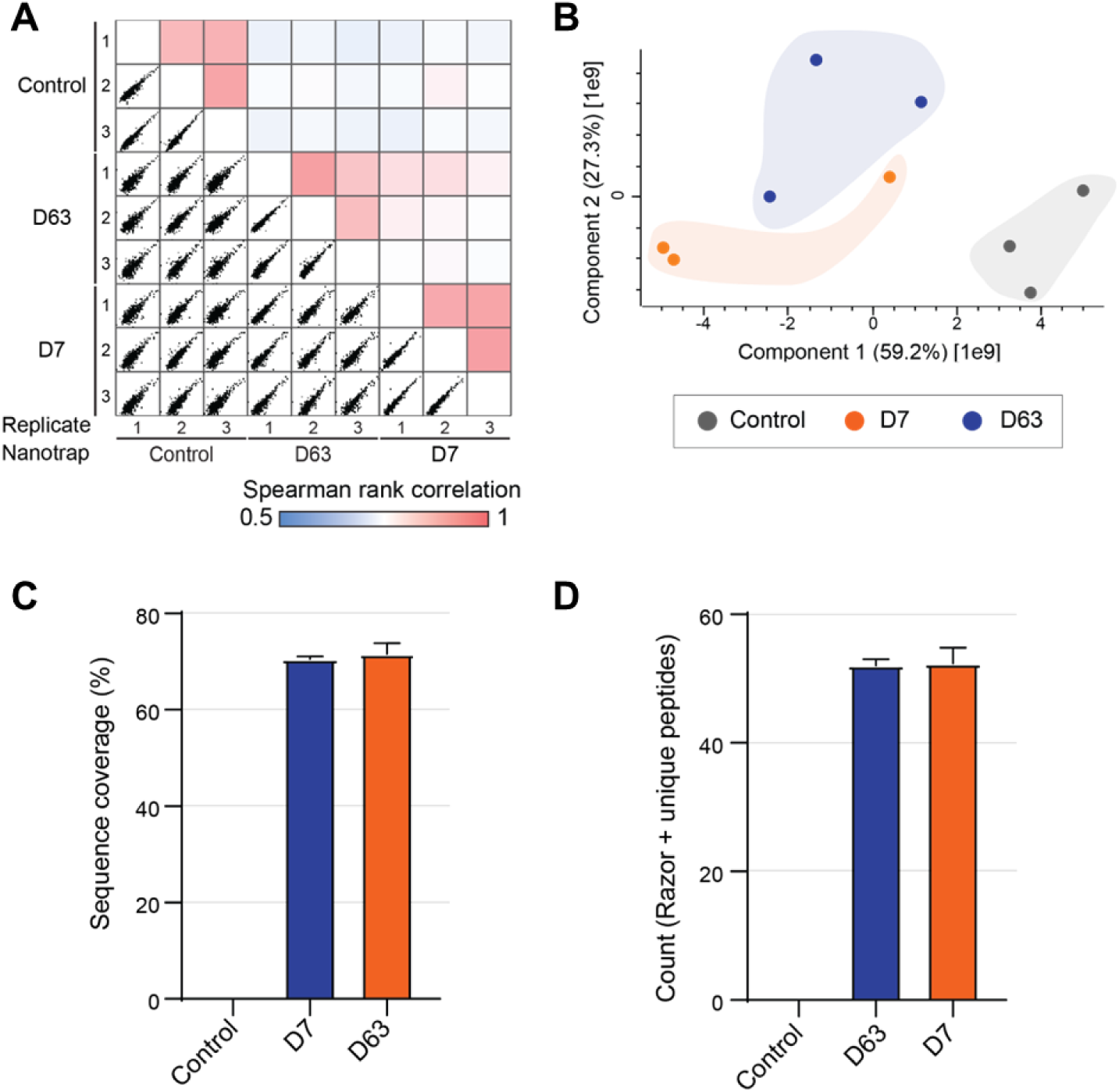
Enrichment efficiency of Drp1-nanotraps. (**A**) Multi Spearman correlation between replicates and nanotraps as indicated. (**B**) Principal component analysis (PCA) reflects the highest similarity between replicates. (**C**) Averaged sequence coverage of Drp1 after precipitation with indicated nanotraps. (**D**) Averaged identification of Drp1 razor and unique peptides for isoform 1, 5 or 7. (**C, D**) Shown are the results of three technical replicates (n=3) ± S.D.

**Supplementary Figure 5:**
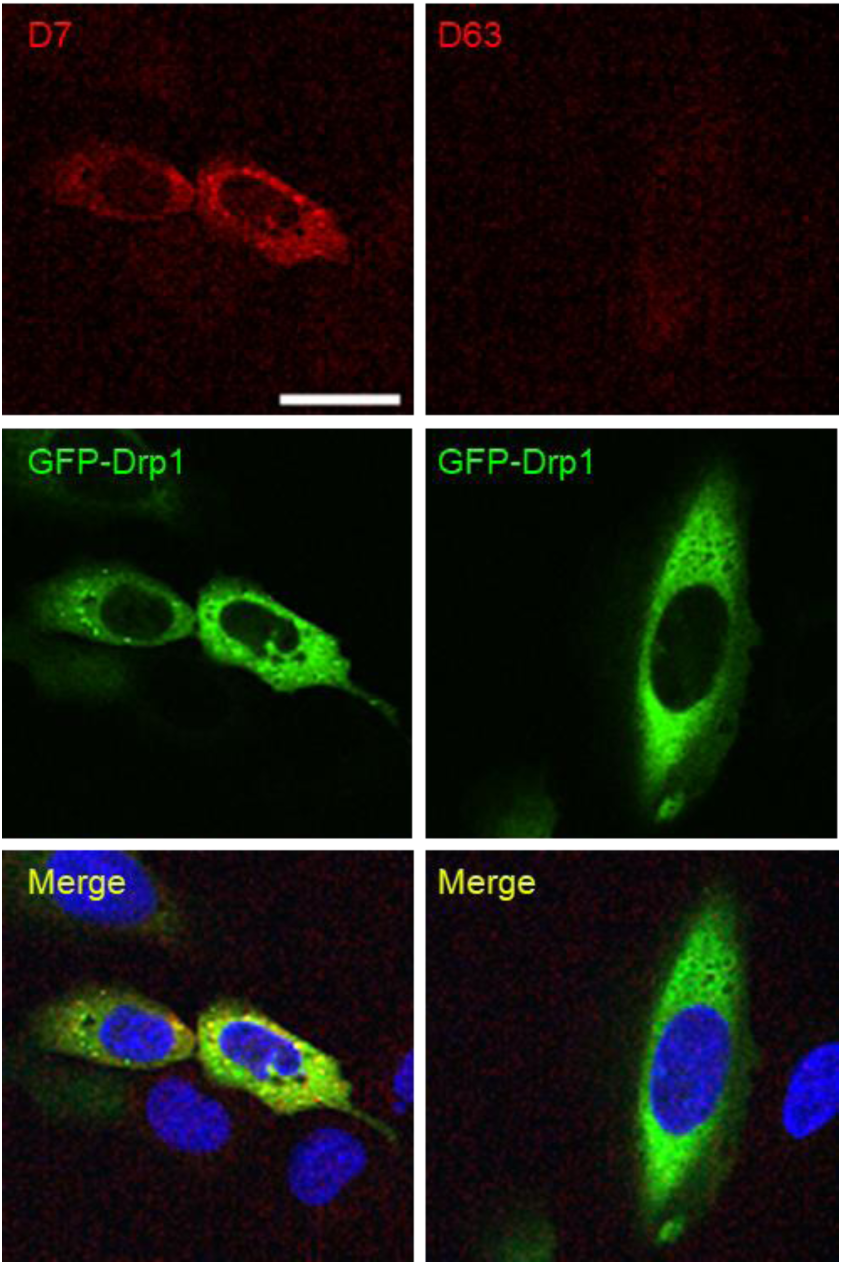
Immunofluorescence staining of GFP-Drp1-expressing cells with Drp1-Nbs. Immunofluorescence (IF) detection of Drp1 in fixed and permeabilized U2OS cells transiently expressing GFP-Drp1 (green) after staining with D7 and D63 (red) as primary labelling probes. Representative confocal laser scanning microscopy (CLSM) images are shown. Scale bar 25 µm.

**Supplementary Figure 6:**
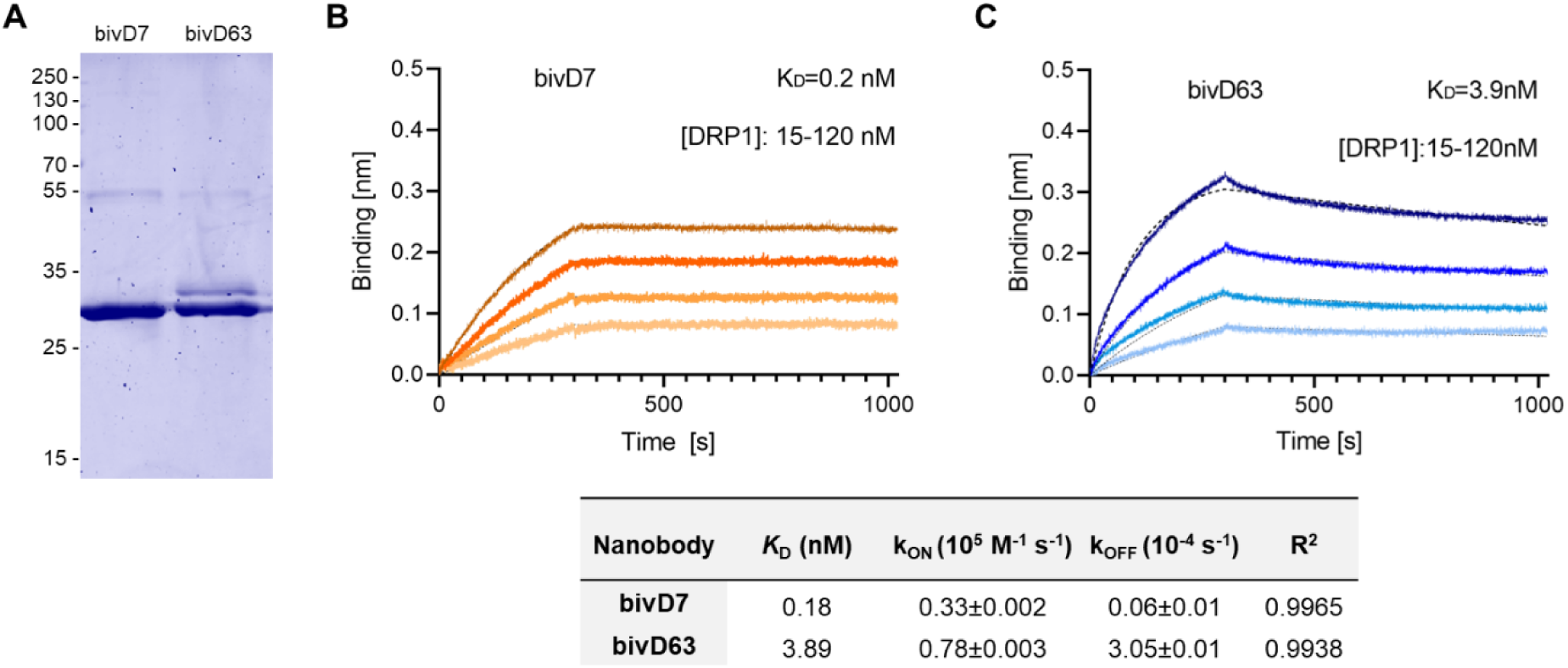
Biochemical characterization of bivalent D7- and D63s. (**A**) Recombinant expression and purification of Drp1-Nbs bivD7 and bivD63 from ExpiCHO cells by immobilized metal affinity chromatography (IMAC). Coomassie staining of purified Nbs (2 µg) is shown. Numbers indicate molecular weight (kDa). (**B, C**) Biolayer interferometry (BLI)-based affinity measurements for bivD7 (**B**) and bivD63 (**C**). Biotinylated Nbs were immobilized on streptavidin sensors. Kinetic measurements were performed using four concentrations of purified Drp1 ranging from 15-120 nM (displayed with gradually darker shades of color). The binding affinity (K_D_) was calculated from global 1:1 fits shown as dashed lines. The table summarizes the affinity analysis of bivD7 and bivD63. Affinities (K_D_), association constants (*k*_on_) and dissociation constants (*k*_off_) are shown as mean ± SD.

**Supplementary Figure 7:**
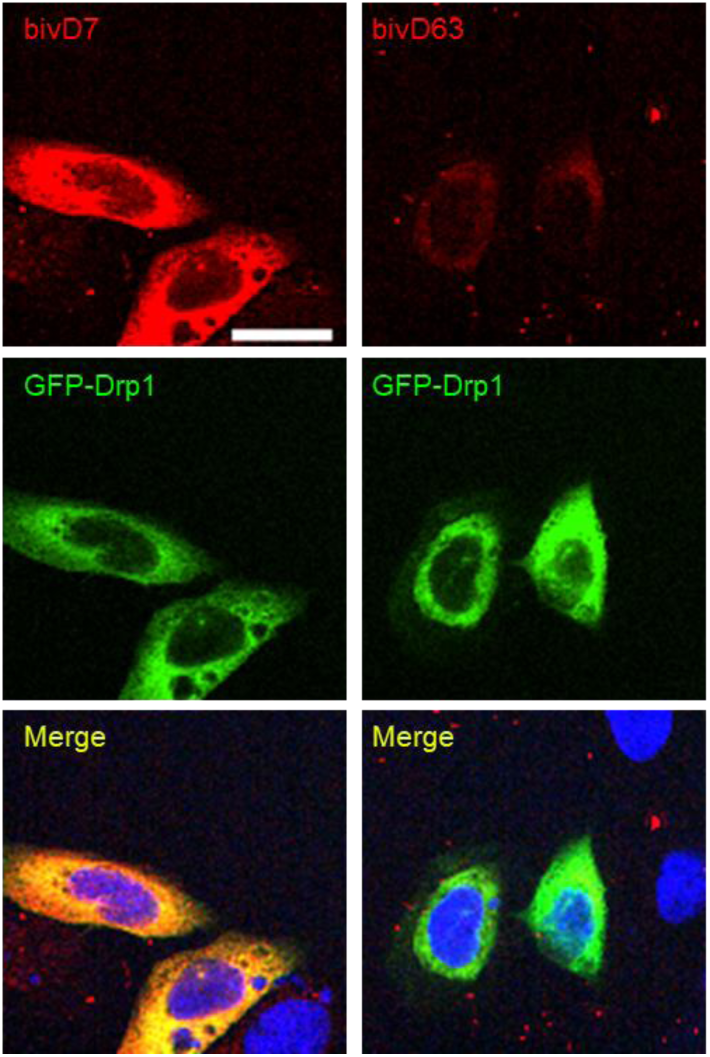
Immunofluorescence staining of GFP-Drp1 expressing cells with bivalent Drp1-Nbs. Immunofluorescence (IF) detection of Drp1 after staining with bivD7 and bivD63 (red) as primary labelling probes in fixed and permeabilized U2OS cells transiently expressing GFP-Drp1 (green). Representative confocal laser scanning microscopy (CLSM) images are shown. Scale bar 25 µm.

**Supplementary Figure 8:**
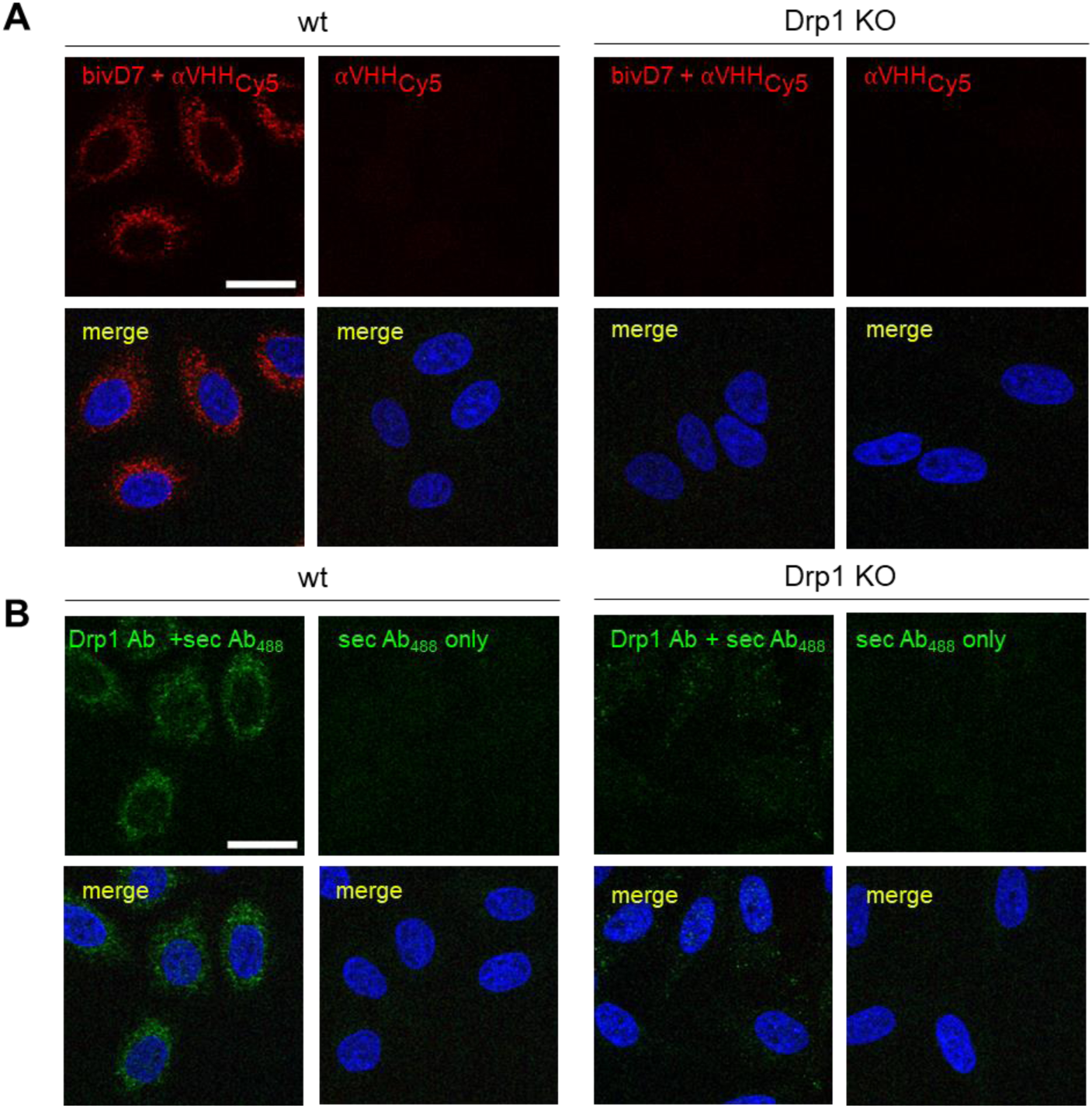
bivD7 specifically recognizes endogenous Drp1. Immunofluorescence (IF) detection of Drp1 in fixed and permeabilised HeLa wt (left panel) or Drp1 KO HeLa cells (right panel) with bivD7 (red) (**A**) or an anti-Drp1 antibody (Drp1 Ab, green) (**B**) as primary labelling probes. For control, cells were stained with the respective fluorescently-labelled secondary antibodies (sec Ab_488_ or αVHH_Cy5_, red) only. Nuclei were counterstained with DAPI (blue). Representative confocal laser scanning microscopy (CLSM) images are shown. Scale bar 25 µm.

**Supplementary Figure 9:**
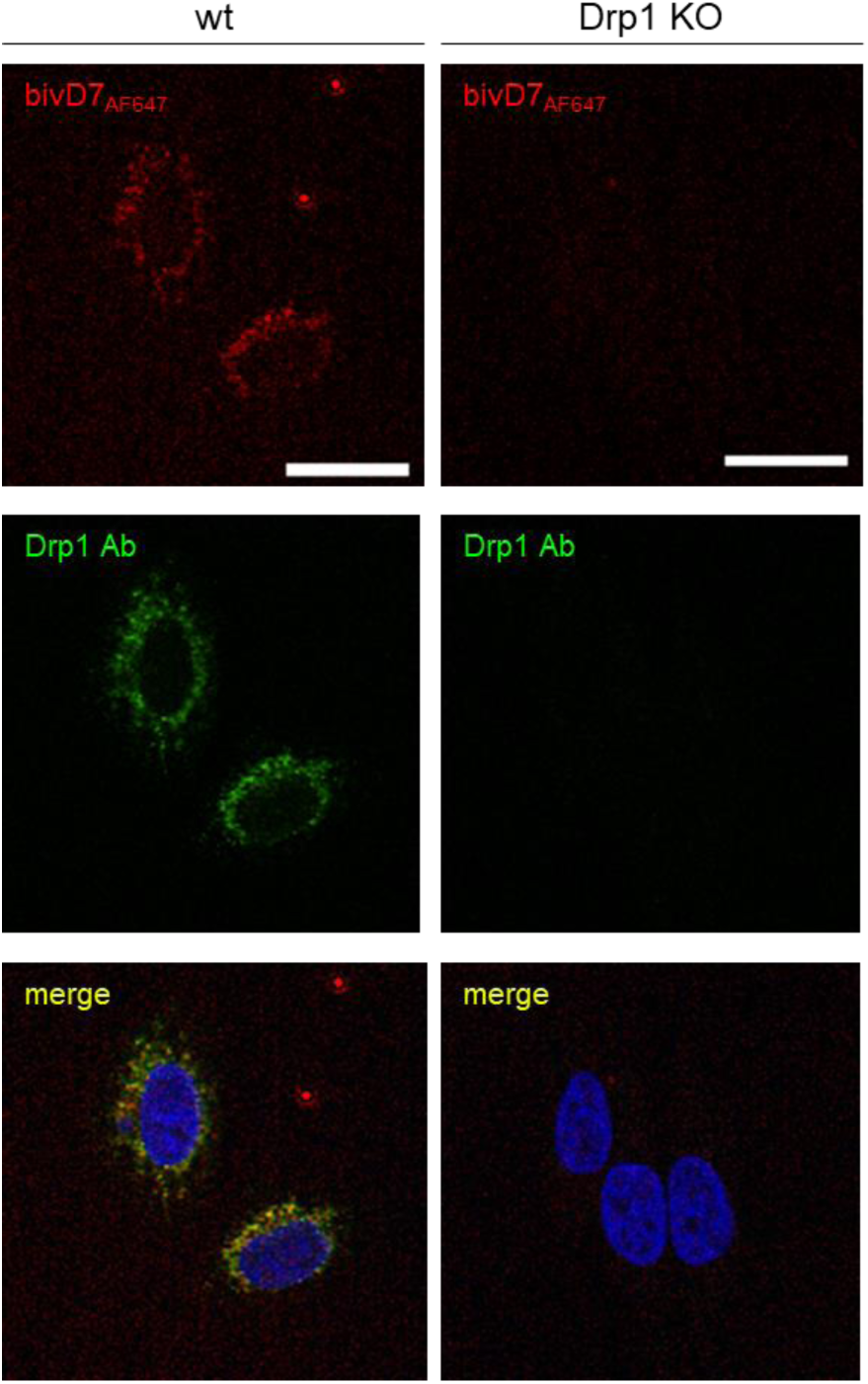
Directly labelled bivD7_AF647_ stained endogenous Drp1. wt or Drp1 KO HeLa cells were stained with fluorescently labelled bivD7_AF647_(red). For control, cells an anti-Drp1 antibody (Drp1 Ab) in combination with an AlexaFluor488-labelled secondary antibody (green) was used. Nuclei were stained with DAPI (blue). Representative confocal laser scanning microscopy (CLSM) images are shown. Scale bar 25 µm.

**Supplementary Figure 10:**
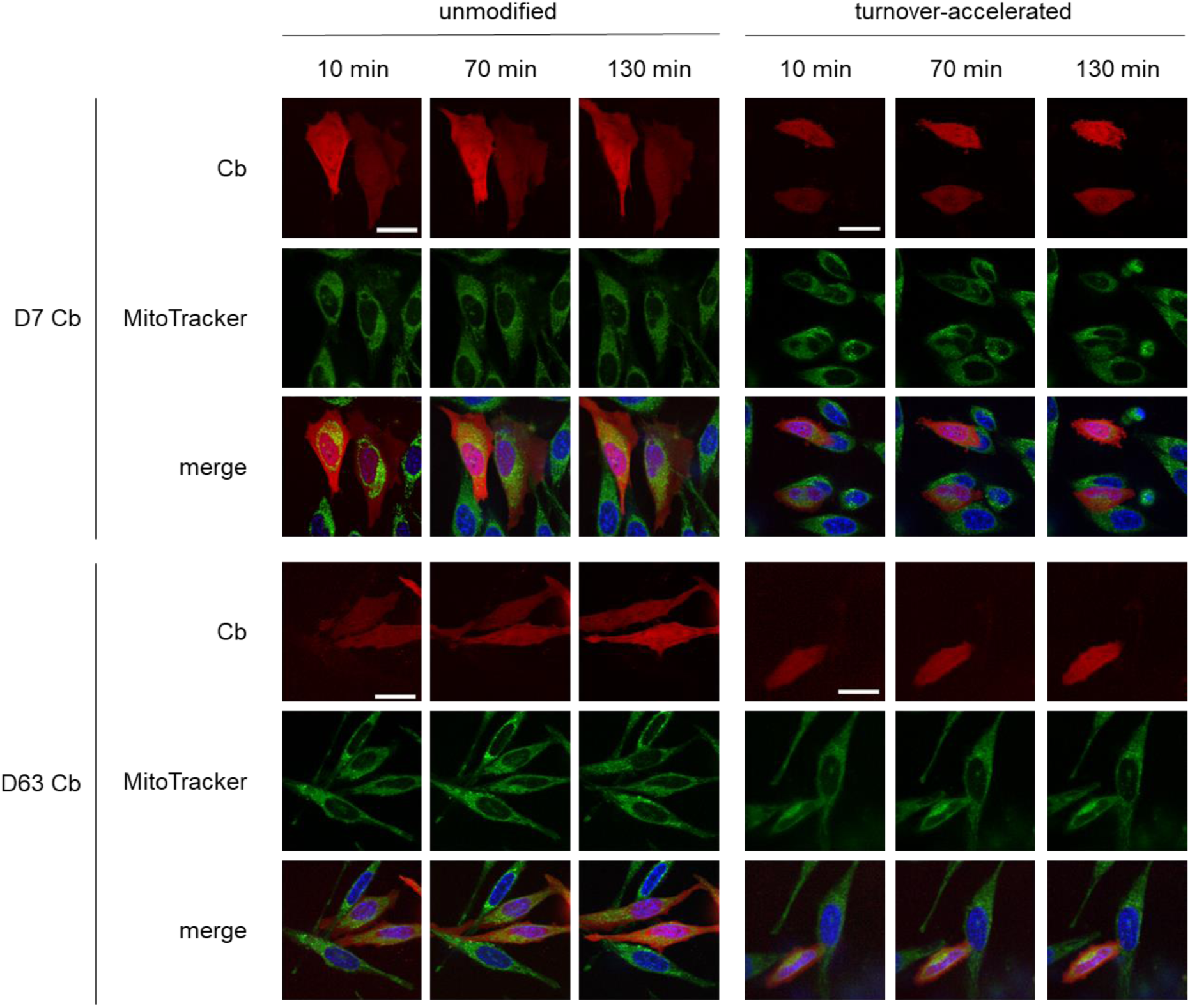
Live Cell imaging of Cbs in Drp1 KO HeLa cells after CCCP treatment. Time lapse microscopy of Drp1 KO HeLa cells transiently expressing unmodified or turnover-accelerated Drp1 Cbs. One day post transfection, cells were stained with Hoechst33258 (blue) and MitoTracker green (green) and subsequently treated with 20 µM CCCP. Representative confocal images were taken after 10 min, 70 min, and 130 min of treatment. Signal intensity was adjusted for each construct to visualize the residual expression of Drp1 Cbs. Scale bar 25 µm.

**Supplementary Figure 11:**
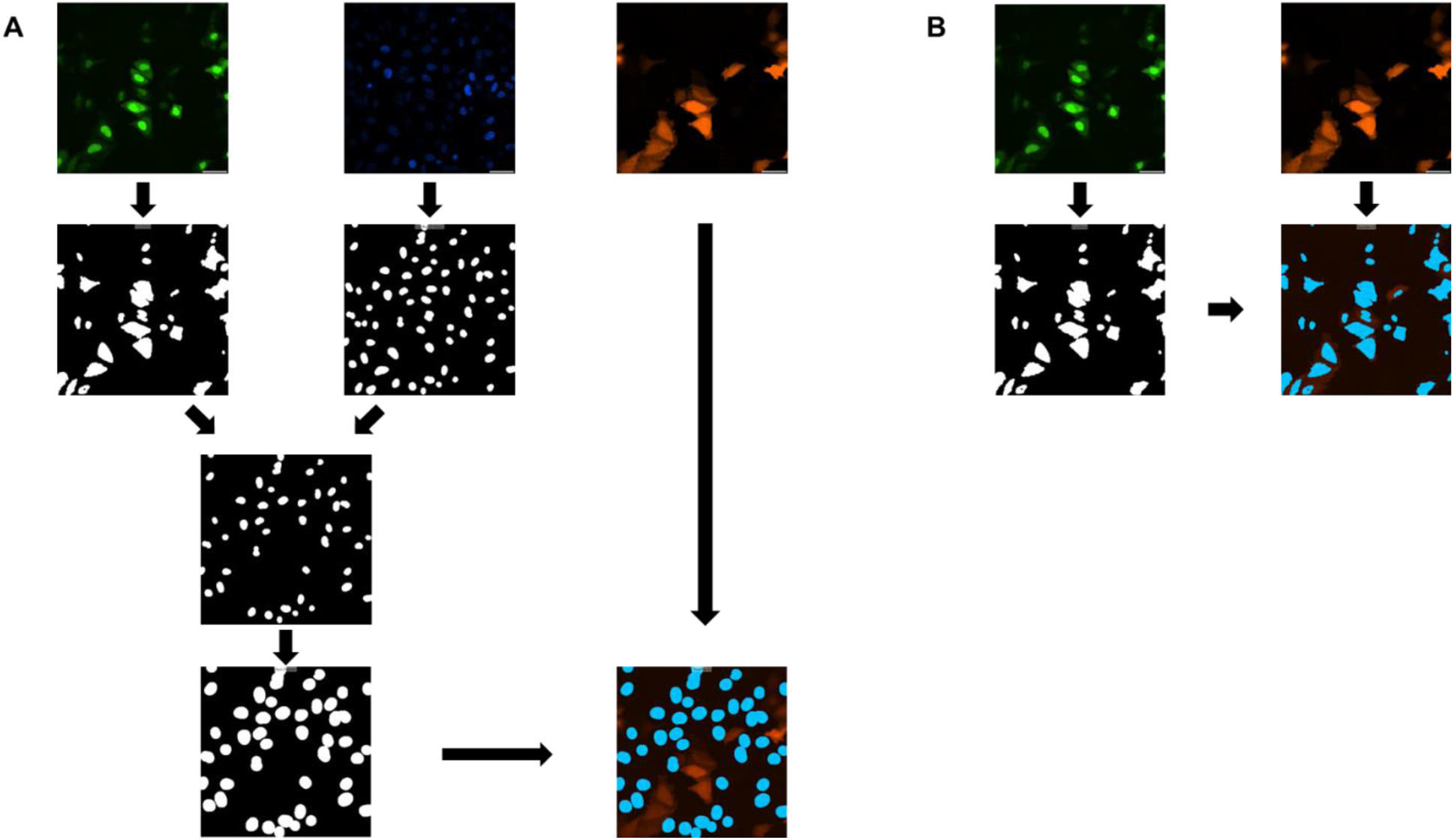
Workflow of the automated image segmentation used for Cb signal quantification. For quantitative image analysis, cells were co-transfected with GFP-NLS(green) and Cb (red), and nuclei were stained using DAPI (blue). (**A**) Workflow for background signal determination. Area of GFP-NLS-expressing cells and nuclei were determined individually (second row). Then, nuclei of GFP-NLS-expressing cells were excluded (third row). Next, the DAPI signal was used to create a cell shape and the RFP intensity was measured in this area (forth row). (**B**) Workflow for Cb signal intensity determination. The GFP signal was used to determine the cell shape and the RFP signal was measured in this area.

**Supplementary Figure 12:**
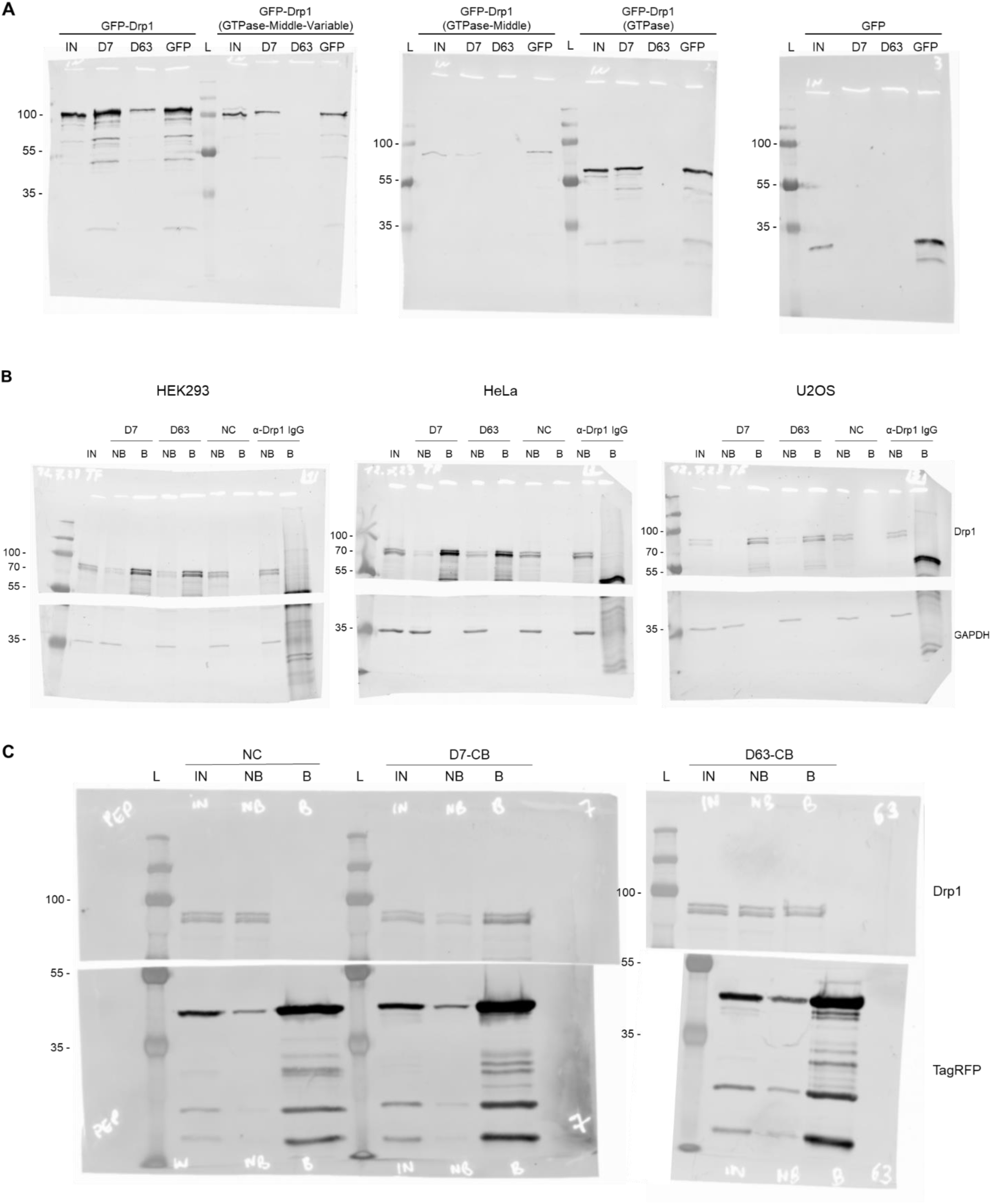
Full-size Western Blots images shown in this study. (**A**) Full size of Western blots depicted in **Fig. 2B** (domain mapping) stained with an anti-GFP antibody. (**B**) Full size of Western blots depicted in **Fig. 3A** and **Fig. S4**. Top half stained with anti-Drp1 antibody; bottom half stained with anti-GAPDH antibody. (**C**) Full size Western blots depicted in **Fig. 5A**. Top half stained with anti-Drp1 antibody; bottom half stained with anti-TagRFP antibody. Molecular weights in kDa are shown on the left of each blot.

### Supplementary Tables

**Supplementary Table 1:**
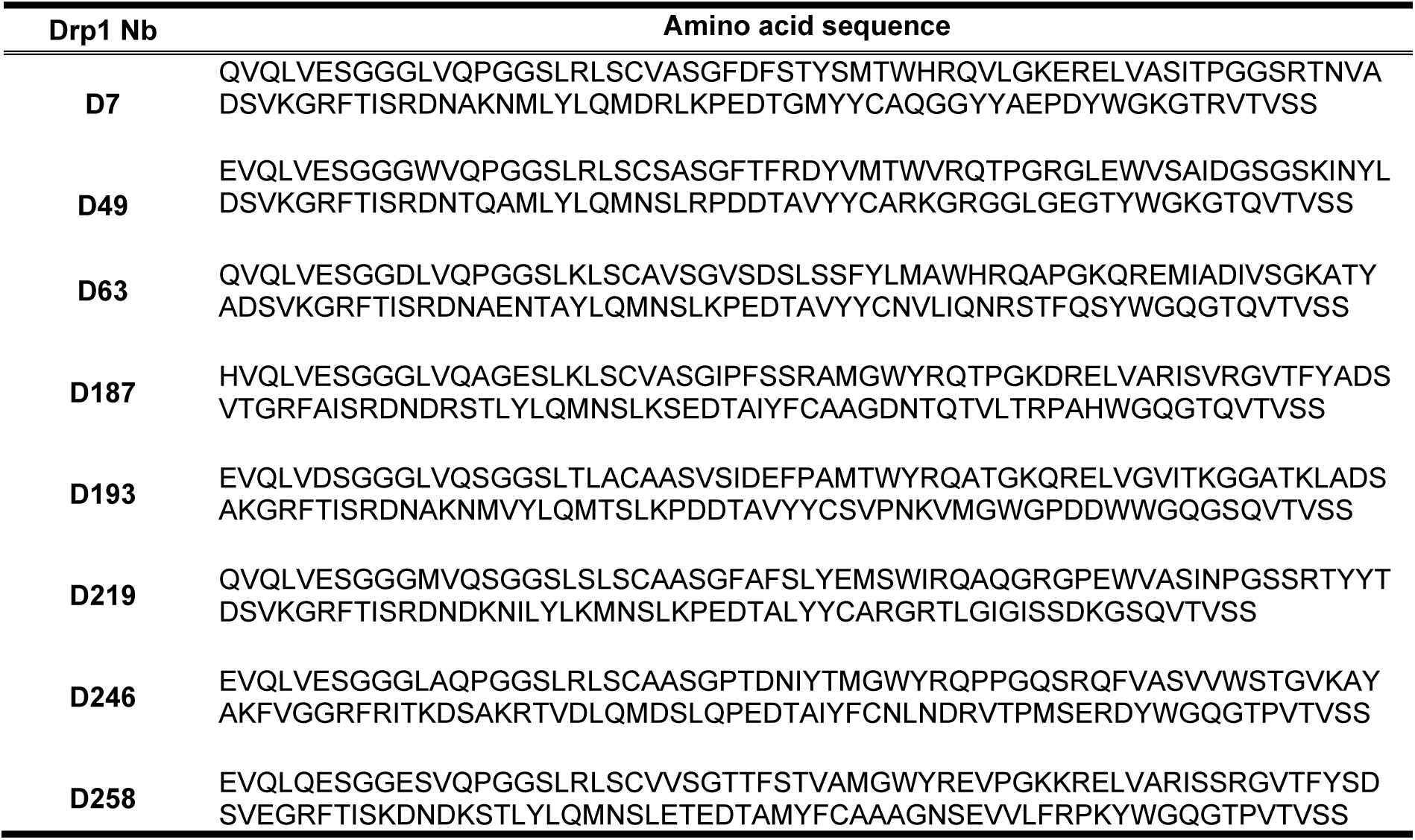
Amino acid sequences of Drp1 Nbs identified as positive clones by Phage ELISA.

**Supplementary Table 2:**
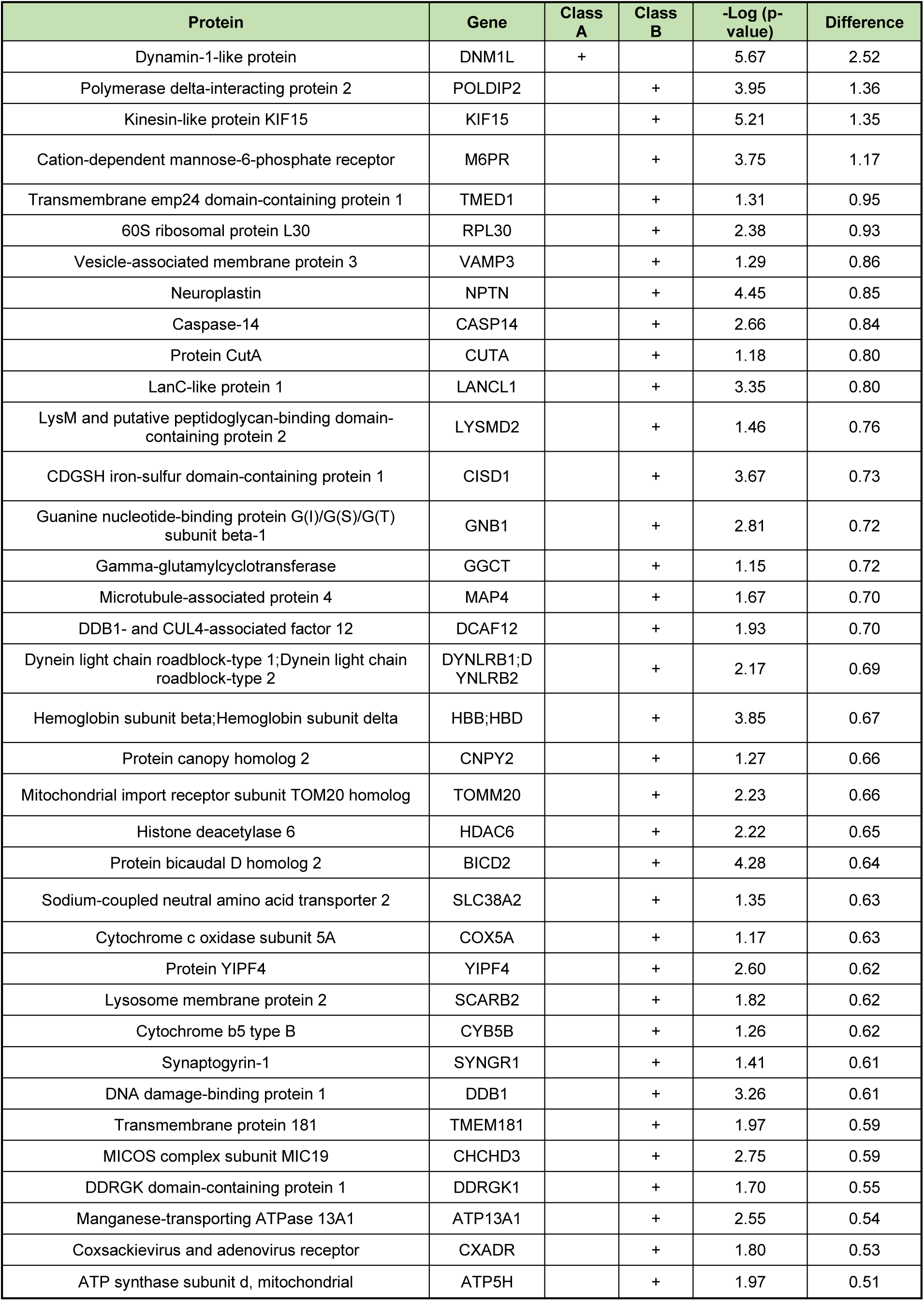
List of significant proteins enriched with the D7 nanotrap. List of class A (FDR=0.01) and class B (FDR=0.05) interactors of Drp1 precipitated by the D7-nanotrap. -Log(p-value) and difference between D7- and control nanotrap are stated for each protein

**Supplementary Table 3:**
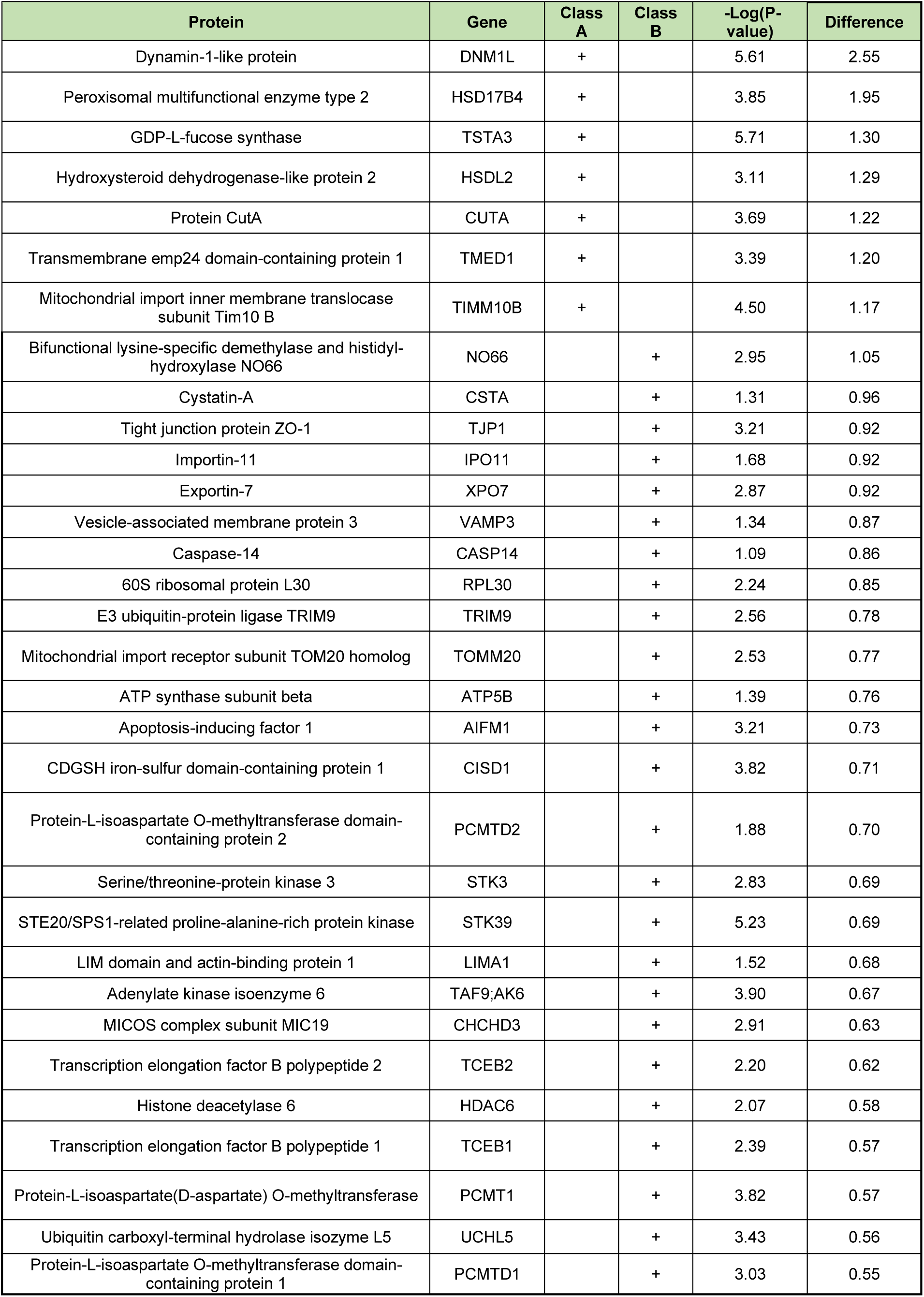
List of significant proteins enriched with the D63 nanotrap. List of class A (FDR=0.01) and class B (FDR=0.05) interactors of Drp1 precipitated by the D63-nanotrap. -Log(p-value) and difference between D63- and control nanotrap are stated for each protein.

**Supplementary Table 4:**
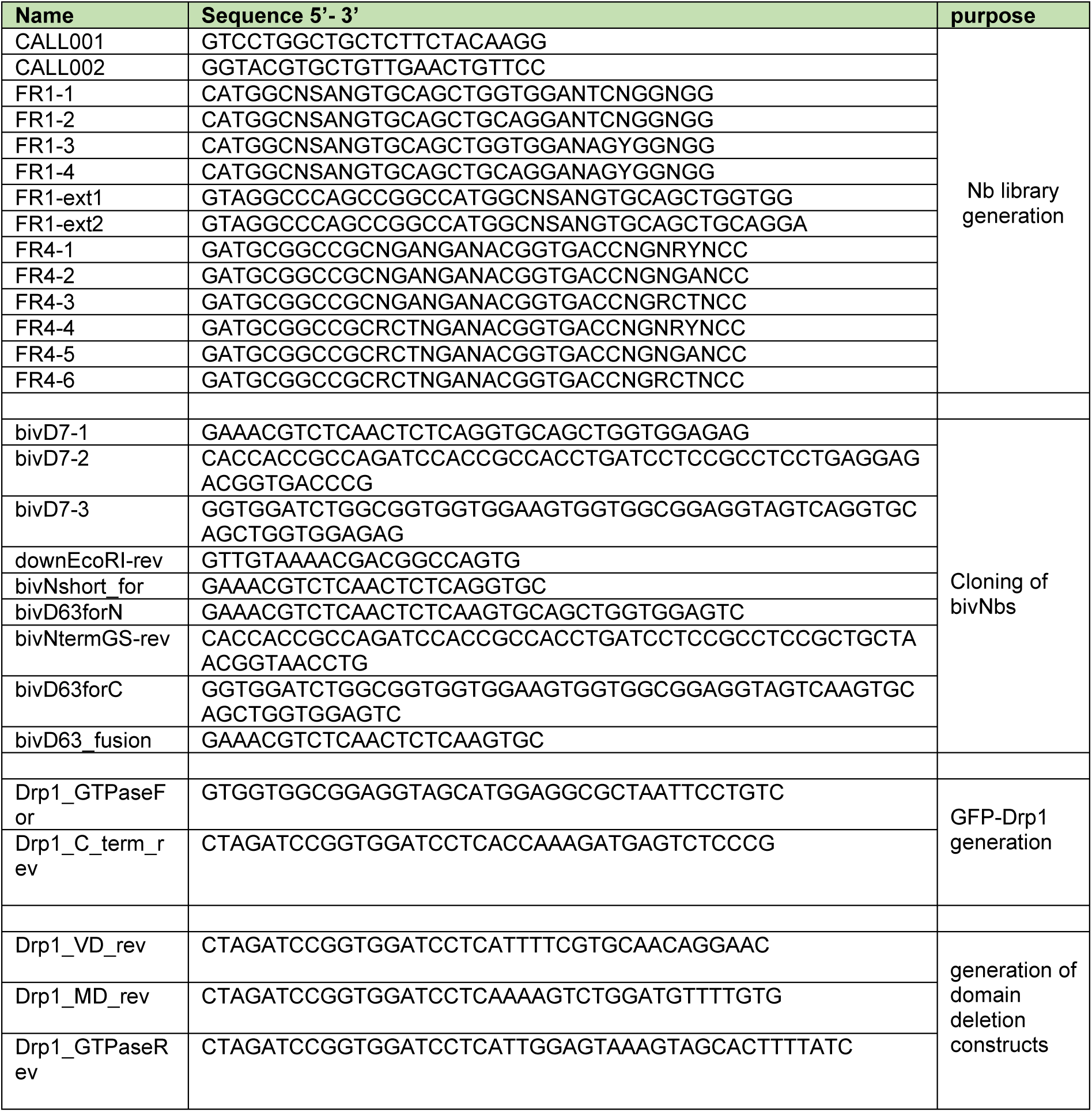
List of oligonucleotides used in this study.

**Supplementary Table 5:** List of expression construct used in this study.

| construct | origin |
| --- | --- |
| pHEN4 | (Arbabi Ghahroudi et al., 1997) |
| pHEN6C | (Rothbauer et al., 2008) |
| D7 in pHEN6C | This study |
| D63 in pHEN6C | This study |
| pCDNA3.4 | Addgene |
| bivD7 in pCDNA3.4 | This study |
| bivD63 in pCDNA3.4 | This study |
| pTagRFP | (Panza et al., 2015) |
| D7 in pTagRFP | This study |
| D63 in pTagRFP | This study |
| pEGFP-Ubi-R-3xGS-tagRFP | (Keller et al., 2018) |
| D7 in pEGFP-Ubi-R-3xGS-tagRFP | This study |
| D63 in pEGFP-Ubi-R-3xGS-tagRFP | This study |
| AcGFP-Drp1 | (Jenner et al., 2022) |
| pEGFP-N1 | Clontech |
| Drp1 in pEGFP-N1 | This study |
| GTPase-MD-VD in pEGFP-N1 | This study |
| GTPase-MD in pEGFP-N1 | This study |
| GTPase in pEGFP-N1 | This study |
| pEGFP-C1 | Clontech |
| Pep-Cb in pTagRFP | (Traenkle et al., 2020) |
| BC2T-GFP-NLS in pEGFPN1 | (Virant et al., 2018) |
| pTYB2 | (Jenner et al., 2022) |

**Supplementary Table 6:** List of antibodies used in this study.

| target | animal | clone | conjugation | manufacturer | catalog number |
| --- | --- | --- | --- | --- | --- |
| Drp1 | rabbit | D6C7 | - | Cell Signaling Technology | #8570 |
| GAPDH | rabbit | polyclonal | - | Invitrogen | PA1-987 |
| GFP | rat | 3H9 | - | ChromoTek | - |
| TagRFP | rabbit | polyclonal | - | Evrogen | AB234 |
| M13 | mouse | monoclonal | HRP | GE Healthcare | 27-9421-01 |
| rabbit | donkey | polyclonal | AF647 | Invitrogen | #A-31573 |
| rabbit | goat | polyclonal | AF488 | Invitrogen | #A-11008 |
| rat | donkey | polyclonal | AF647 | Invitrogen | #A-21247 |
| alpaca | goat | polyclonal | Cy5 | Jackson Immunoresearch | 128-175-232 |

## References

Arbabi Ghahroudi, M., A. Desmyter, L. Wyns, R. Hamers, and S. Muyldermans. 1997. Selection and identification of single domain antibody fragments from camel heavy-chain antibodies. FEBS Lett. 414:521–526.

Bai, J., Y. Lei, G.L. An, and L. He. 2015. Down-regulation of deacetylase HDAC6 inhibits the melanoma cell line A375.S2 growth through ROS-dependent mitochondrial pathway. PLoS One. 10:e0121247.

Bolte, S., and F.P. Cordelières. 2006. A guided tour into subcellular colocalization analysis in light microscopy. J Microsc. 224:213–232.

Braun, M.B., B. Traenkle, P.A. Koch, F. Emele, F. Weiss, O. Poetz, T. Stehle, and U. Rothbauer. 2016. Peptides in headlock--a novel high-affinity and versatile peptide-binding nanobody for proteomics and microscopy. Sci Rep. 6:19211.

Burgess, A., T. Lorca, and A. Castro. 2012. Quantitative live imaging of endogenous DNA replication in mammalian cells. PLoS One. 7:e45726.

Burgstaller, S., T.R. Wagner, H. Bischof, S. Bueckle, A. Padamsey, D. Frecot, P.D. Kaiser, D. Skrabak, R. Malli, R. Lukowski, and U. Rothbauer. 2022. Monitoring extracellular ion and metabolite dynamics with recombinant nanobody-fused biosensors. iScience. 25:104907.

Carrington, G., D. Tomlinson, and M. Peckham. 2019. Exploiting nanobodies and Affimers for superresolution imaging in light microscopy. Mol Biol Cell. 30:2737–2740.

Chakrabarti, R., W.K. Ji, R.V. Stan, J. de Juan Sanz, T.A. Ryan, and H.N. Higgs. 2018. INF2-mediated actin polymerization at the ER stimulates mitochondrial calcium uptake, inner membrane constriction, and division. J Cell Biol. 217:251–268.

Chen, C.H., S.L. Howng, S.L. Hwang, C.K. Chou, C.H. Liao, and Y.R. Hong. 2000. Differential expression of four human dynamin-like protein variants in brain tumors. DNA Cell Biol. 19:189–194.

Cho, D.H., T. Nakamura, J. Fang, P. Cieplak, A. Godzik, Z. Gu, and S.A. Lipton. 2009. S-nitrosylation of Drp1 mediates beta-amyloid-related mitochondrial fission and neuronal injury. Science. 324:102–105.

Cox, J., and M. Mann. 2008. MaxQuant enables high peptide identification rates, individualized p.p.b.-range mass accuracies and proteome-wide protein quantification. Nat Biotechnol. 26:1367–1372.

Cramer, K., A.L. Bolender, I. Stockmar, R. Jungmann, R. Kasper, and J.Y. Shin. 2019. Visualization of Bacterial Protein Complexes Labeled with Fluorescent Proteins and Nanobody Binders for STED Microscopy. Int J Mol Sci. 20.

De Vos, K.J., V.J. Allan, A.J. Grierson, and M.P. Sheetz. 2005. Mitochondrial function and actin regulate dynamin-related protein 1-dependent mitochondrial fission. Curr Biol. 15:678–683.

Driouchi, A., S.D. Gray-Owen, and C.M. Yip. 2022. Correlated STORM-homoFRET imaging reveals highly heterogeneous membrane receptor structures. J Biol Chem. 298:102448.

English, K., and M.C. Barton. 2021. HDAC6: A Key Link Between Mitochondria and Development of Peripheral Neuropathy. Front Mol Neurosci. 14:684714.

Fagbadebo, F.O., P.D. Kaiser, K. Zittlau, N. Bartlick, T.R. Wagner, T. Froehlich, G. Jarjour, S. Nueske, A. Scholz, B. Traenkle, B. Macek, and U. Rothbauer. 2022. A Nanobody-Based Toolset to Monitor and Modify the Mitochondrial GTPase Miro1. Front Mol Biosci. 9:835302.

Ferreira-da-Silva, A., C. Valacca, E. Rios, H. Pópulo, P. Soares, M. Sobrinho-Simões, L. Scorrano, V. Máximo, and S. Campello. 2015. Mitochondrial dynamics protein Drp1 is overexpressed in oncocytic thyroid tumors and regulates cancer cell migration. PLoS One. 10:e0122308.

Frecot, D.I., T. Froehlich, and U. Rothbauer. 2023. 30 years of nanobodies - an ongoing success story of small binders in biological research. J Cell Sci. 136.

Friedman, J.R., L.L. Lackner, M. West, J.R. DiBenedetto, J. Nunnari, and G.K. Voeltz. 2011. ER tubules mark sites of mitochondrial division. Science. 334:358–362.

Frohlich, C., S. Grabiger, D. Schwefel, K. Faelber, E. Rosenbaum, J. Mears, O. Rocks, and O. Daumke. 2013. Structural insights into oligomerization and mitochondrial remodelling of dynamin 1-like protein. EMBO J. 32:1280–1292.

Fruh, S.M., U. Matti, P.R. Spycher, M. Rubini, S. Lickert, T. Schlichthaerle, R. Jungmann, V. Vogel, J. Ries, and I. Schoen. 2021. Site-Specifically-Labeled Antibodies for Super-Resolution Microscopy Reveal In Situ Linkage Errors. ACS Nano. 15:12161–12170.

Giacomello, M., A. Pyakurel, C. Glytsou, and L. Scorrano. 2020. The cell biology of mitochondrial membrane dynamics. Nat Rev Mol Cell Biol. 21:204–224.

Götzke, H., M. Kilisch, M. Martínez-Carranza, S. Sograte-Idrissi, A. Rajavel, T. Schlichthaerle, N. Engels, R. Jungmann, P. Stenmark, F. Opazo, and S. Frey. 2019. The ALFA-tag is a highly versatile tool for nanobody-based bioscience applications. Nat Commun. 10:4403.

Gross, G.G., J.A. Junge, R.J. Mora, H.B. Kwon, C.A. Olson, T.T. Takahashi, E.R. Liman, G.C. Ellis-Davies, A.W. McGee, B.L. Sabatini, R.W. Roberts, and D.B. Arnold. 2013. Recombinant probes for visualizing endogenous synaptic proteins in living neurons. Neuron. 78:971–985.

Hamers-Casterman, C., T. Atarhouch, S. Muyldermans, G. Robinson, C. Hamers, E.B. Songa, N. Bendahman, and R. Hamers. 1993. Naturally occurring antibodies devoid of light chains. Nature. 363:446–448.

Han, H., J. Tan, R. Wang, H. Wan, Y. He, X. Yan, J. Guo, Q. Gao, J. Li, S. Shang, F. Chen, R. Tian, W. Liu, L. Liao, B. Tang, and Z. Zhang. 2020. PINK1 phosphorylates Drp1(S616) to regulate mitophagy-independent mitochondrial dynamics. EMBO Rep. 21:e48686.

Hua, J., Z. Gao, S. Zhong, B. Wei, J. Zhu, and R. Ying. 2021. CISD1 protects against atherosclerosis by suppressing lipid accumulation and inflammation via mediating Drp1. Biochem Biophys Res Commun. 577:80–88.

Huang, T.L., C.R. Chang, C.Y. Chien, G.K. Huang, Y.F. Chen, L.J. Su, H.T. Tsai, Y.S. Lin, F.M. Fang, and C.H. Chen. 2022. DRP1 contributes to head and neck cancer progression and induces glycolysis through modulated FOXM1/MMP12 axis. Mol Oncol. 16:2585–2606.

Ingerman, E., and J. Nunnari. 2005. A continuous, regenerative coupled GTPase assay for dynamin-related proteins. Methods Enzymol. 404:611–619.

Itoh, K., Y. Adachi, T. Yamada, T.L. Suzuki, T. Otomo, H.M. McBride, T. Yoshimori, M. Iijima, and H. Sesaki. 2018. A brain-enriched Drp1 isoform associates with lysosomes, late endosomes, and the plasma membrane. J Biol Chem. 293:11809–11822.

Iwata, R., P. Casimir, and P. Vanderhaeghen. 2020. Mitochondrial dynamics in postmitotic cells regulate neurogenesis. Science. 369:858–862.

Jenner, A., A. Pena-Blanco, R. Salvador-Gallego, B. Ugarte-Uribe, C. Zollo, T. Ganief, J. Bierlmeier, M. Mund, J.E. Lee, J. Ries, D. Schwarzer, B. Macek, and A.J. Garcia-Saez. 2022. DRP1 interacts directly with BAX to induce its activation and apoptosis. EMBO J. 41:e108587.

Ji, W.K., R. Chakrabarti, X. Fan, L. Schoenfeld, S. Strack, and H.N. Higgs. 2017. Receptor-mediated Drp1 oligomerization on endoplasmic reticulum. J Cell Biol. 216:4123–4139.

Jin, J.Y., X.X. Wei, X.L. Zhi, X.H. Wang, and D. Meng. 2021. Drp1-dependent mitochondrial fission in cardiovascular disease. Acta Pharmacol Sin. 42:655–664.

Kashatus, J.A., A. Nascimento, L.J. Myers, A. Sher, F.L. Byrne, K.L. Hoehn, C.M. Counter, and D.F. Kashatus. 2015. Erk2 phosphorylation of Drp1 promotes mitochondrial fission and MAPK-driven tumor growth. Mol Cell. 57:537–551.

Keller, B.M., J. Maier, K.A. Secker, S.M. Egetemaier, Y. Parfyonova, U. Rothbauer, and B. Traenkle. 2018. Chromobodies to Quantify Changes of Endogenous Protein Concentration in Living Cells. Mol Cell Proteomics. 17:2518–2533.

Kim, E.Y., Y. Zhang, I. Beketaev, A.M. Segura, W. Yu, Y. Xi, J. Chang, and J. Wang. 2015. SENP5, a SUMO isopeptidase, induces apoptosis and cardiomyopathy. J Mol Cell Cardiol. 78:154–164.

Koch, A., M. Thiemann, M. Grabenbauer, Y. Yoon, M.A. McNiven, and M. Schrader. 2003. Dynamin-like protein 1 is involved in peroxisomal fission. J Biol Chem. 278:8597–8605.

Koch, J., and C. Brocard. 2012. PEX11 proteins attract Mff and human Fis1 to coordinate peroxisomal fission. J Cell Sci. 125:3813–3826.

Korobova, F., V. Ramabhadran, and H.N. Higgs. 2013. An actin-dependent step in mitochondrial fission mediated by the ER-associated formin INF2. Science. 339:464–467.

Labrousse, A.M., M.D. Zappaterra, D.A. Rube, and A.M. van der Bliek. 1999. C. elegans dynamin-related protein DRP-1 controls severing of the mitochondrial outer membrane. Mol Cell. 4:815–826.

Li, S., S. Xu, B.A. Roelofs, L. Boyman, W.J. Lederer, H. Sesaki, and M. Karbowski. 2015. Transient assembly of F-actin on the outer mitochondrial membrane contributes to mitochondrial fission. J Cell Biol. 208:109–123.

Li, X., and S.J. Gould. 2002. PEX11 promotes peroxisome division independently of peroxisome metabolism. J Cell Biol. 156:643–651.

Liang, J., Y. Yang, L. Bai, F. Li, and E. Li. 2020. DRP1 upregulation promotes pancreatic cancer growth and metastasis through increased aerobic glycolysis. J Gastroenterol Hepatol. 35:885–895.

Loson, O.C., Z. Song, H. Chen, and D.C. Chan. 2013. Fis1, Mff, MiD49, and MiD51 mediate Drp1 recruitment in mitochondrial fission. Mol Biol Cell. 24:659–667.

Maidorn, M., A. Olichon, S.O. Rizzoli, and F. Opazo. 2019. Nanobodies reveal an extra-synaptic population of SNAP-25 and Syntaxin 1A in hippocampal neurons. MAbs. 11:305–321.

Maier, J., B. Traenkle, and U. Rothbauer. 2015. Real-time analysis of epithelial-mesenchymal transition using fluorescent single-domain antibodies. Sci Rep. 5:13402.

Michalska, B.M., K. Kwapiszewska, J. Szczepanowska, T. Kalwarczyk, P. Patalas-Krawczyk, K. Szczepański, R. Hołyst, J. Duszyński, and J. Szymański. 2018. Insight into the fission mechanism by quantitative characterization of Drp1 protein distribution in the living cell. Sci Rep. 8:8122.

Montecinos-Franjola, F., B.L. Bauer, J.A. Mears, and R. Ramachandran. 2020. GFP fluorescence tagging alters dynamin-related protein 1 oligomerization dynamics and creates disassembly-refractory puncta to mediate mitochondrial fission. Sci Rep. 10:14777.

Nan, J., W. Zhu, M.S. Rahman, M. Liu, D. Li, S. Su, N. Zhang, X. Hu, H. Yu, M.P. Gupta, and J. Wang. 2017. Molecular regulation of mitochondrial dynamics in cardiac disease. Biochim Biophys Acta Mol Cell Res. 1864:1260–1273.

Osellame, L.D., A.P. Singh, D.A. Stroud, C.S. Palmer, D. Stojanovski, R. Ramachandran, and M.T. Ryan. 2016. Cooperative and independent roles of the Drp1 adaptors Mff, MiD49 and MiD51 in mitochondrial fission. J Cell Sci. 129:2170–2181.

Otera, H., N. Ishihara, and K. Mihara. 2013. New insights into the function and regulation of mitochondrial fission. Biochim Biophys Acta. 1833:1256–1268.

Ovesny, M., P. Krizek, J. Borkovec, Z. Svindrych, and G.M. Hagen. 2014. ThunderSTORM: a comprehensive ImageJ plug-in for PALM and STORM data analysis and super-resolution imaging. Bioinformatics. 30:2389–2390.

Palmer, C.S., K.D. Elgass, R.G. Parton, L.D. Osellame, D. Stojanovski, and M.T. Ryan. 2013. Adaptor proteins MiD49 and MiD51 can act independently of Mff and Fis1 in Drp1 recruitment and are specific for mitochondrial fission. J Biol Chem. 288:27584–27593.

Panza, P., J. Maier, C. Schmees, U. Rothbauer, and C. Söllner. 2015. Live imaging of endogenous protein dynamics in zebrafish using chromobodies. Development. 142:1879–1884.

Pardon, E., T. Laeremans, S. Triest, S.G. Rasmussen, A. Wohlkönig, A. Ruf, S. Muyldermans, W.G. Hol, B.K. Kobilka, and J. Steyaert. 2014. A general protocol for the generation of Nanobodies for structural biology. Nat Protoc. 9:674–693.

Popp, M.W., and H.L. Ploegh. 2011. Making and breaking peptide bonds: protein engineering using sortase. Angew Chem Int Ed Engl. 50:5024–5032.

Rasmussen, M.L., N. Taneja, A.C. Neininger, L. Wang, G.L. Robertson, S.N. Riffle, L. Shi, B.C. Knollmann, D.T. Burnette, and V. Gama. 2020. MCL-1 Inhibition by Selective BH3 Mimetics Disrupts Mitochondrial Dynamics Causing Loss of Viability and Functionality of Human Cardiomyocytes. iScience. 23:101015.

Rehman, J., H.J. Zhang, P.T. Toth, Y. Zhang, G. Marsboom, Z. Hong, R. Salgia, A.N. Husain, C. Wietholt, and S.L. Archer. 2012. Inhibition of mitochondrial fission prevents cell cycle progression in lung cancer. Faseb j. 26:2175–2186.

Rothbauer, U., K. Zolghadr, S. Muyldermans, A. Schepers, M.C. Cardoso, and H. Leonhardt. 2008. A versatile nanotrap for biochemical and functional studies with fluorescent fusion proteins. Mol Cell Proteomics. 7:282–289.

Rothbauer, U., K. Zolghadr, S. Tillib, D. Nowak, L. Schermelleh, A. Gahl, N. Backmann, K. Conrath, S. Muyldermans, M.C. Cardoso, and H. Leonhardt. 2006. Targeting and tracing antigens in live cells with fluorescent nanobodies. Nat Methods. 3:887–889.

Schindelin, J., I. Arganda-Carreras, E. Frise, V. Kaynig, M. Longair, T. Pietzsch, S. Preibisch, C. Rueden, S. Saalfeld, B. Schmid, J.Y. Tinevez, D.J. White, V. Hartenstein, K. Eliceiri, P. Tomancak, and A. Cardona. 2012. Fiji: an open-source platform for biological-image analysis. Nat Methods. 9:676–682.

Serasinghe, M.N., S.Y. Wieder, T.T. Renault, R. Elkholi, J.J. Asciolla, J.L. Yao, O. Jabado, K. Hoehn, Y. Kageyama, H. Sesaki, and J.E. Chipuk. 2015. Mitochondrial division is requisite to RAS-induced transformation and targeted by oncogenic MAPK pathway inhibitors. Mol Cell. 57:521–536.

Sharp, W.W., Y.H. Fang, M. Han, H.J. Zhang, Z. Hong, A. Banathy, E. Morrow, J.J. Ryan, and S.L. Archer. 2014. Dynamin-related protein 1 (Drp1)-mediated diastolic dysfunction in myocardial ischemia-reperfusion injury: therapeutic benefits of Drp1 inhibition to reduce mitochondrial fission. Faseb j. 28:316–326.

Shevchenko, A., H. Tomas, J. Havlis, J.V. Olsen, and M. Mann. 2006. In-gel digestion for mass spectrometric characterization of proteins and proteomes. Nat Protoc. 1:2856–2860.

Shirendeb, U.P., M.J. Calkins, M. Manczak, V. Anekonda, B. Dufour, J.L. McBride, P. Mao, and P.H. Reddy. 2012. Mutant huntingtin’s interaction with mitochondrial protein Drp1 impairs mitochondrial biogenesis and causes defective axonal transport and synaptic degeneration in Huntington’s disease. Hum Mol Genet. 21:406–420.

Sibler, A.P., J. Courtete, C.D. Muller, G. Zeder-Lutz, and E. Weiss. 2005. Extended half-life upon binding of destabilized intrabodies allows specific detection of antigen in mammalian cells. FEBS J. 272:2878–2891.

Solesio, M.E., S. Saez-Atienzar, J. Jordan, and M.F. Galindo. 2013. 3-Nitropropionic acid induces autophagy by forming mitochondrial permeability transition pores rather than activating the mitochondrial fission pathway. Br J Pharmacol. 168:63–75.

Song, W., J. Chen, A. Petrilli, G. Liot, E. Klinglmayr, Y. Zhou, P. Poquiz, J. Tjong, M.A. Pouladi, M.R. Hayden, E. Masliah, M. Ellisman, I. Rouiller, R. Schwarzenbacher, B. Bossy, G. Perkins, and E. Bossy-Wetzel. 2011. Mutant huntingtin binds the mitochondrial fission GTPase dynamin-related protein-1 and increases its enzymatic activity. Nat Med. 17:377–382.

Tang, J.C., E. Drokhlyansky, B. Etemad, S. Rudolph, B. Guo, S. Wang, E.G. Ellis, J.Z. Li, and C.L. Cepko. 2016. Detection and manipulation of live antigen-expressing cells using conditionally stable nanobodies. Elife. 5.

Tong, M., D. Zablocki, and J. Sadoshima. 2020. The role of Drp1 in mitophagy and cell death in the heart. J Mol Cell Cardiol. 142:138–145.

Traenkle, B., F. Emele, R. Anton, O. Poetz, R.S. Haeussler, J. Maier, P.D. Kaiser, A.M. Scholz, S. Nueske, A. Buchfellner, T. Romer, and U. Rothbauer. 2015. Monitoring interactions and dynamics of endogenous beta-catenin with intracellular nanobodies in living cells. Mol Cell Proteomics. 14:707–723.

Traenkle, B., S. Segan, F.O. Fagbadebo, P.D. Kaiser, and U. Rothbauer. 2020. A novel epitope tagging system to visualize and monitor antigens in live cells with chromobodies. Sci Rep. 10:14267.

Vantaggiato, C., M. Castelli, M. Giovarelli, G. Orso, M.T. Bassi, E. Clementi, and C. De Palma. 2019. The Fine Tuning of Drp1-Dependent Mitochondrial Remodeling and Autophagy Controls Neuronal Differentiation. Front Cell Neurosci. 13:120.

Varshavsky, A. 2005. Ubiquitin fusion technique and related methods. Methods Enzymol. 399:777–799.

Virant, D., B. Traenkle, J. Maier, P.D. Kaiser, M. Bodenhöfer, C. Schmees, I. Vojnovic, B. Pisak-Lukáts, U. Endesfelder, and U. Rothbauer. 2018. A peptide tag-specific nanobody enables high-quality labeling for dSTORM imaging. Nat Commun. 9:930.

Wagner, T.R., E. Ostertag, P.D. Kaiser, M. Gramlich, N. Ruetalo, D. Junker, J. Haering, B. Traenkle, M. Becker, A. Dulovic, H. Schweizer, S. Nueske, A. Scholz, A. Zeck, K. Schenke-Layland, A. Nelde, M. Strengert, J.S. Walz, G. Zocher, T. Stehle, M. Schindler, N. Schneiderhan-Marra, and U. Rothbauer. 2021. NeutrobodyPlex-monitoring SARS-CoV-2 neutralizing immune responses using nanobodies. EMBO Rep. 22:e52325.

Wagner, T.R., and U. Rothbauer. 2020. Nanobodies Right in the Middle: Intrabodies as Toolbox to Visualize and Modulate Antigens in the Living Cell. Biomolecules. 10.

Wang, X., B. Su, H.G. Lee, X. Li, G. Perry, M.A. Smith, and X. Zhu. 2009. Impaired balance of mitochondrial fission and fusion in Alzheimer’s disease. J Neurosci. 29:9090–9103.

Wegner, W., P. Ilgen, C. Gregor, J. van Dort, A.C. Mott, H. Steffens, and K.I. Willig. 2017. In vivo mouse and live cell STED microscopy of neuronal actin plasticity using far-red emitting fluorescent proteins. Sci Rep. 7:11781.

Xie, L., F. Shi, Y. Li, W. Li, X. Yu, L. Zhao, M. Zhou, J. Hu, X. Luo, M. Tang, J. Fan, J. Zhou, Q. Gao, W. Wu, X. Zhang, W. Liao, A.M. Bode, and Y. Cao. 2020. Drp1-dependent remodeling of mitochondrial morphology triggered by EBV-LMP1 increases cisplatin resistance. Signal Transduct Target Ther. 5:56.

Xiong, X., S. Hasani, L.E.A. Young, D.R. Rivas, A.T. Skaggs, R. Martinez, C. Wang, H.L. Weiss, M.S. Gentry, R.C. Sun, and T. Gao. 2022. Activation of Drp1 promotes fatty acids-induced metabolic reprograming to potentiate Wnt signaling in colon cancer. Cell Death Differ. 29:1913–1927.

Yu, R., S.B. Jin, M. Ankarcrona, U. Lendahl, M. Nistér, and J. Zhao. 2021. The Molecular Assembly State of Drp1 Controls its Association With the Mitochondrial Recruitment Receptors Mff and MIEF1/2. Front Cell Dev Biol. 9:706687.

Yu, R., T. Liu, C. Ning, F. Tan, S.B. Jin, U. Lendahl, J. Zhao, and M. Nistér. 2019. The phosphorylation status of Ser-637 in dynamin-related protein 1 (Drp1) does not determine Drp1 recruitment to mitochondria. J Biol Chem. 294:17262–17277.

Zerihun, M., S. Sukumaran, and N. Qvit. 2023. The Drp1-Mediated Mitochondrial Fission Protein Interactome as an Emerging Core Player in Mitochondrial Dynamics and Cardiovascular Disease Therapy. Int J Mol Sci. 24.

Zhao, J., U. Lendahl, and M. Nistér. 2013. Regulation of mitochondrial dynamics: convergences and divergences between yeast and vertebrates. Cell Mol Life Sci. 70:951–976.

Zhu, P.P., A. Patterson, J. Stadler, D.P. Seeburg, M. Sheng, and C. Blackstone. 2004. Intra- and intermolecular domain interactions of the C-terminal GTPase effector domain of the multimeric dynamin-like GTPase Drp1. J Biol Chem. 279:35967–35974.

Zunino, R., A. Schauss, P. Rippstein, M. Andrade-Navarro, and H.M. McBride. 2007. The SUMO protease SENP5 is required to maintain mitochondrial morphology and function. J Cell Sci. 120:1178–1188.

